# Trex-QTL: A mixture-model for identification of genetic effects with global effects on molecular phenotypes

**DOI:** 10.64898/2026.08.07.743622

**Authors:** Cynthia Wu, Andrey Bzikadze, Tianyao Xu, Eric M. Mendenhall, Hao Chen, Francesca Telese, Oksana Polesskaya, Daniel Munro, Abraham A. Palmer, Alon Goren, Melissa Gymrek

## Abstract

While thousands of *cis* expression quantitative trait loci (*cis*-eQTLs) have been reliably identified, detecting *trans*-eQTL effects has proven to be challenging due to insufficient statistical power, lack of comparable tissues and cohorts, and low reproducibility across studies. Here, we present Trex-QTL, a novel *trans*-eQTL detection method that models eQTL summary statistics as a mixture consisting of both target gene and null associations. Compared to other recently developed methods, Trex-QTL has improved power for *trans*-eQTL detection and employs a simplified framework, requiring only eQTL association summary statistics as input. We performed extensive simulations to characterize the conditions under which *trans*-eQTLs are detectable by Trex-QTL across a range of effect sizes and numbers of target genes. We applied Trex-QTL to the Depression Genes and Networks (DGN) dataset and replicated two well-established *trans*-eQTLs at *ARHGEF3* and *IKZF1*. We then applied Trex-QTL to the deeply characterized heterogeneous stock (HS) rat cohort with matched brain transcriptomic and genomic data, identifying 7 top-scoring, linkage disequilibrium-independent *trans*-eQTLs. One previously unreported *trans*-eQTL is at the locus harboring *Jag2*, a critical ligand for the Notch signaling pathway, which is associated with decreased *Jag2* expression and decreased expression of multiple downstream genes including known Notch targets. A second example is a strong *trans*-eQTL overlapping a cluster of interferon genes associated with interferon-response genes including *C4a* and *Parp14*. We show evidence that this signal is mediated by a *cis*-eQTL for a cluster of interferon ligand genes that operate upstream of interferon receptor signaling. Both of these signals co-localize with association signals for a range of other phenotypes measured in this cohort. Overall, we demonstrate that Trex-QTL represents a powerful method to identify *trans*-eQTLs with global effects on molecular phenotypes and identify novel biologically compelling examples of such loci.

## Introduction

Genetic variants that are associated with changes in gene expression, termed eQTLs, are major drivers of complex traits and human disease^1^. The majority of eQTL studies have focused on *cis*-eQTLs, for which the genetic variant is near the target gene. Genetic variants that are not close to the target gene (*trans*-eQTLs) are also thought to play an important role in gene regulation and disease risk^2,3,4^, but have been far more challenging to analyze. Specifically, *trans*-eQTL identification is typically underpowered due to the large number of putative gene-by-variant pairs resulting in a high multiple testing burden and reduced statistical power. Additionally, *trans* effects are generally weaker than *cis* effects^5^, and therefore require larger sample sizes for reliable detection. Further, technical sources of noise including batch effects, sequencing and alignment artifacts, and other latent confounders, can introduce substantial variation in expression datasets and lead to many false positive eQTL calls^6^. Thus, most identified *trans*-eQTLs have not been consistently replicated across studies due to the insufficient statistical power, lack of comparable tissues and cohorts, and potential false positive associations^2^.

Previous studies applied various methods to detect *trans*-eQTLs. Albert and Bloom, *et al.*^7^ detected *trans*-eQTLs clustering at 102 hotspot loci in yeast segregants by exhaustively testing putative gene-by-variant pairs. However in species with larger genomes, such as humans, pairwise testing becomes impractical due to the large number of tests that are required. To improve power, alternative strategies aggregate expression signals across sets of genes. Kolberg, *et al.*^8^ tested associations between variants and co-expression-derived gene modules, identifying replicable *trans*-eQTLs in human blood, though results depended strongly on the choice of co-expression method. Brynedal, *et al.*^9^ leveraged cross-phenotype meta-analysis (CPMA)^10^ to detect variants with global *trans* effects by evaluating whether their association statistics across genes deviate from the null distribution. Yet, this approach is best suited for detecting *trans*-eQTLs influencing many genes and has limited power to detect *trans*-eQTLs with a small number of target genes. More recently, Trans-PCO^11^ applied a principal component-based omnibus test to combine expression signals within predefined gene modules. However, this approach requires gene modules to be specified a priori, and thus the resulting associations are dependent on the choice of module definition. Another method, ARCHIE^12^ leverages summary statistics, linkage disequilibrium, and gene co-expression to identify trait-specific *trans*-association patterns, but depends on high-quality external reference datasets to obtain co-expression structure. PLIER^13^ uses matrix factorization guided by prior pathway annotations to extract biologically interpretable latent variables, but relies on the quality of prior gene sets and is not tailored for variant-level *trans*-eQTL detection. A related approach, MultiXcan^14^ integrates predicted gene expression across tissues to improve gene-trait association detection, but depends on accurate expression prediction models and does not directly model variant effects across all genes.

Here, we introduce **Trex-QTL** (*Trans* Regulatory Effect eXplorer for QTLs), a novel *trans*-eQTL detection method that improves power over pairwise methods by jointly modeling the effects of an individual variant across all genes. While Trex-QTL aggregates information across genes in a similar manner to CPMA, trans-PCO, and ARCHIE, it differs by employing a biologically motivated mixture model of effect sizes. Another advantage of Trex-QTL is that it relies on summary statistics alone and does not require specification of dimensionality reduction parameters or definition of gene modules. We developed a simulation framework to benchmark Trex-QTL against established approaches (CPMA and pairwise analysis). We first applied Trex-QTL to the Depression Genes and Networks^15^ (DGN) dataset, consisting of whole-blood RNA-sequencing and genotypes from 922 individuals, and replicated well-established *trans*-eQTLs. We then leveraged a rich dataset from 339 deeply phenotyped outbred rats with matched brain hemisphere transcriptomes and whole-genome sequencing to identify novel *trans*-eQTLs, including loci associated with Notch signaling targets and interferon-related genes.

## Results

### Trex-QTL identifies trans-QTLs with global effects

We developed Trex-QTL, a method for detecting genetic variants with global effects in *trans* on a molecular phenotype of interest (**Fig. 1a**). The Trex-QTL framework is agnostic to the molecular phenotype being tested, but we focus on the application of detecting variants with global effects on gene expression (*trans*-eQTLs) to illustrate the method. The premise of Trex-QTL is that the *P*-values of pairwise association statistics between a variant of interest and expression of each gene are drawn from two distinct distributions: one for the target genes and another for the non-target genes, in line with the biological expectation that only a subset of the genes are impacted. Under the null hypothesis that the variant is not a *trans*-eQTL, all (-log) *P*-values are from non-target genes and are expected to follow a standard (λ = 1) exponential distribution. On the other hand, if the variant is a *trans*-eQTL, the *P*-values are drawn from a mixture of two distinct exponential distributions, with one corresponding to non-null associations (λ > 1). Trex-QTL fits this mixture distribution to learn the proportion (*t*) of all genes that are targets (non-null) for each variant and outputs a likelihood ratio statistic (*S*) which can be used to rank candidate *trans*-eQTLs. Finally, it obtains the statistical significance of each candidate based on comparison of the *S* statistic to an empirical null distribution derived from permutation testing (**Methods**).

**Figure 1:**
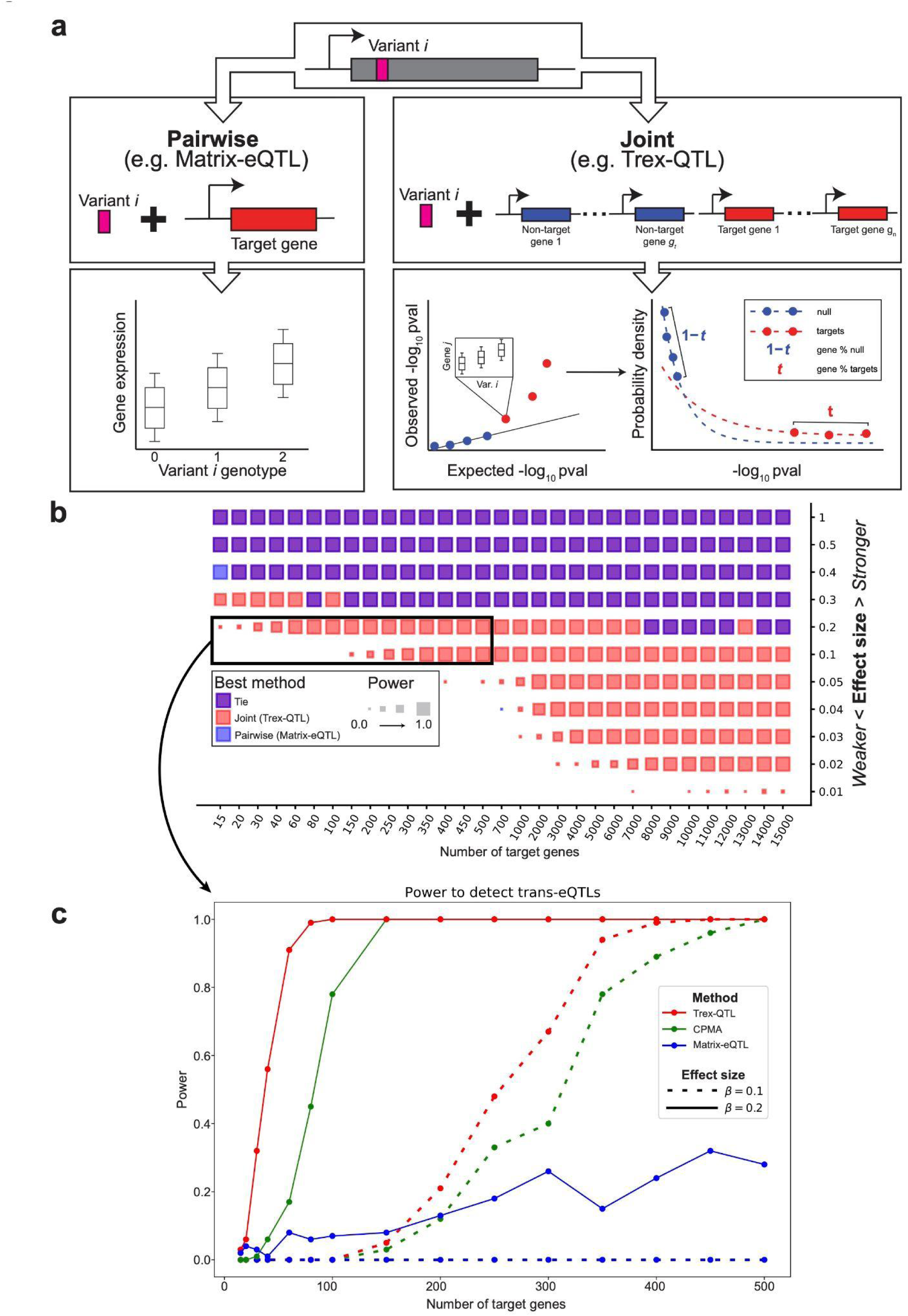
Trex-QTL detects variants with global effects on molecular phenotypes. **a.** *Trans*-eQTLs can affect expression of one or more target genes. We focus on two classes of *trans*-eQTL detection methods. <u>Left:</u> pairwise methods test all possible variant-gene pairs; <u>Right:</u> joint methods perform a single test for each variant by considering all association statistics for that variant together. Trex-QTL along with other methods (e.g. CPMA^9^) are joint methods. Trex-QTL uniquely assumes association statistics can be approximated by a mixture distribution consisting of two distributions, one for the target genes and another for the null (non-target) genes. **b.** Results from our simulation framework represented as a heatmap comparing Trex-QTL vs Matrix-eQTL. The heatmap shows the *trans*-eQTL detection method with highest power for detecting *trans*-eQTLs with varying number of target genes and effect sizes. The color represents the method with highest power: blue=Matrix-eQTL, red=Trex-QTL, purple=tie between Trex-QTL and Matrix-eQTL. Trex-QTL has increased power over Matrix-eQTL to detect *trans*-eQTLs with small number of target genes and effect size. A similar analysis including CPMA is shown in **Supplementary** Fig. 1. **c.** The power of Trex-QTL, CPMA, and Matrix-eQTL is shown for effect sizes (*β*) of 0.1 (weak; dashed lines), or 0.2 (strong; solid lines).

To evaluate Trex-QTL, we developed a framework to simulate expression datasets driven by *trans*-eQTLs with a range of effect sizes and number of target genes for a specified sample size and total number of genes (**Methods**). Our framework further models gene-gene correlation and can simulate effects of unknown technical covariates such as those captured by PEER factors^6^. We used our framework to evaluate the power of Trex-QTL to detect *trans*-eQTLs with various properties (**Fig. 1b-c**) under a setting of 15,000 genes based on the typical number of expressed protein-coding genes in human or other mammalian model organism datasets. For comparison, we additionally evaluated power using CPMA^9^ (which similarly jointly models all association statistics for a particular variant but does not model target vs. non-target genes) as well as the traditional method of performing pairwise tests for each variant-phenotype pair.

We first examined a baseline case without modeling gene-gene correlation or technical covariates. As expected, all methods show increasing power to detect *trans*-eQTLs as a function of effect size and sample size (**Fig. 1b**). For downstream analyses, we focus on results for 500 simulated individuals, similar to sample sizes for available expression datasets^1,15,16^. At this sample size, naive pairwise methods are underpowered to detect all but the strongest effects, whereas joint methods are best for detecting variants affecting 100 or more target genes, with at least modest effect sizes. In cases when the proportion of target genes (*t*) and/or effect size (*β*) are very large (e.g. β>0.2 and *t*>∼1%; **Fig. 1c**), both Trex-QTL and CPMA are able to detect nearly all simulated *trans*-eQTLs. However, Trex-QTL outperforms CPMA for cases where the number of target genes and/or the effect size are modest (**Fig. 1c**; **Supplementary Fig. 1**). We additionally evaluated Trex-QTL’s inferred *t* (proportion of target genes) and observed that in scenarios where Trex-QTL has sufficient power (*t*>∼1% of target genes) the inferred *t* generally aligns with the simulated values across a range of effect sizes, although estimates are modestly inflated in various settings (**Supplementary Fig. 2**).

We then conducted another round of simulations, modeling extensive gene-gene correlation as has been observed in real expression datasets^2^. In this case, we found that *P*-values based on a theoretical null distribution that assumes each pairwise variant-gene test is independent, are severely inflated. On the other hand, *P*-values obtained using a permutation-based null distribution that accounts for gene-gene correlation, are well-calibrated (**Supplementary Fig. 3**). We therefore rely on permutation testing for obtaining *P*-values in downstream analyses.

Finally, we evaluated the impact of the commonly applied correction of including top expression PCs (or similarly, PEER factors^6^) as covariates when computing gene-variant summary statistics upstream of Trex-QTL. For this, we simulated the effects of strong *trans*-eQTLs with or without effects of technical variation as typically modeled by PEER factors and applied Trex-QTL with or without including the top 20 expression PCs as covariates (**Supplementary Fig. 4**). We found that in the absence of simulated technical variation, top expression PCs often capture strong *trans*-eQTLs with global effects, and thus regressing them out substantially reduces detection power. On the other hand, in the presence of simulated technical variation failing to regress out top expression PCs resulted in severe inflation of summary statistics which can lead to false positive calls. Thus, in downstream analyses we include expression PCs as covariates but recognize this comes at the cost of likely excluding some bona-fide *trans*-eQTL signals if their effects are relatively strong compared to those of true technical sources of variation. Even when including top PCs as covariates, Trex-QTL association statistics are still moderately inflated (**Supplementary Fig. 5**), further motivating our choice to use an empirical null distribution based on permutation testing.

### Replication of known strong trans-eQTLs in human whole blood

We used the well studied Depression Genes and Networks^15^ (DGN) dataset to benchmark Trex-QTL. The DGN dataset consists of genotype and whole blood RNA-seq data from 922 individuals of European ancestry. This dataset was designed to study the genetic and transcriptomic properties of depression and other complex traits. The DGN dataset has been widely used for eQTL mapping and has enabled the discovery of at least two strong and replicable *trans*-eQTLs in which genetic variation at a single locus is associated with changes in expression of a large number of target genes. These include: (1) *trans*-eQTL variants in *ARHGEF3*, which were identified by TransPCO^11^, PLIER^13^, and MultiXcan^14^. Further, strong *trans*-eQTL signals at this locus have been replicated in multiple other platelet-specific datasets^8,17^; and (2) *trans*-eQTL variants in *IKZF1* which were identified in DGN by transPCO^11^ and GBAT^18^, as well as in an independent peripheral blood dataset^4^ and a separate whole blood dataset^19^.

Following preprocessing steps (**Methods**), including removal of genes and variants in poorly mapped regions, filtering variants with low minor allele frequency (MAF; <0.05), and pruning variant pairs in high linkage disequilibrium (LD; r^2^ >= 0.95), we retained 11,641 protein coding genes and 1,755,712 variants for downstream analysis. We utilized all covariates already defined by DGN which include genotype and expression PCs, and various biological and technical factors. To focus on *trans*-eQTLs, we regressed out effects of *cis*-eQTLs within 1Mb of each gene from the expression data (**Methods**). This step removes variation in expression driven by *cis* variants, which can often have strong effects on expression of individual genes, and therefore improves power for *trans*-eQTL detection. We then applied Trex-QTL to the *cis*-adjusted expression matrix to identify 93 candidate *trans*-eQTLs (FDR < 25%). The top-ranked signal mapped to the *ARHGEF3* locus, and the second strongest signal mapped to *IKZF1*, replicating the previously reported *trans*-eQTLs in this dataset (**Fig. 2b**). These results demonstrate that Trex-QTL reliably recovers established *trans*-eQTLs in well-characterized cohorts using summary statistics alone. Full trex-QTL summary statistics for the DGN analysis are provided in **Supplementary Dataset 1**.

**Figure 2:**
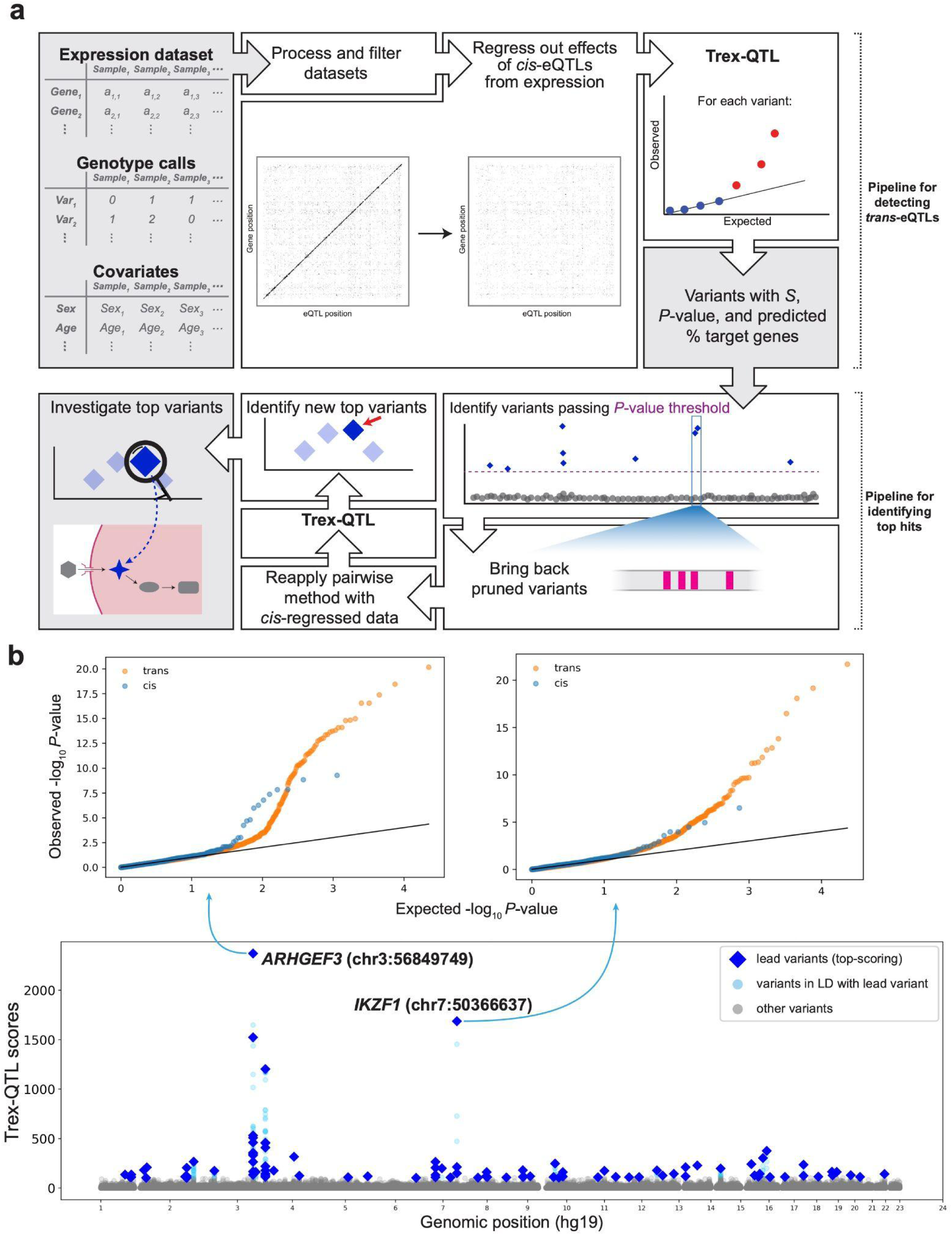
Overview of the Trex-QTL analysis pipeline and application to DGN dataset. **a.** Workflow for detecting *trans*-eQTLs using Trex-QTL. Starting from gene expression, genotype, and covariate data, datasets are first processed and filtered. Effects of *cis*-eQTLs are regressed out from gene expression to reduce confounding from local regulatory signals. The Trex-QTL method is then applied to identify variants with global (*trans*) effects by aggregating association statistics across genes, yielding a score (*S*), permutation *P*-value, and an estimated proportion of affected target genes. Variants passing a significance threshold are retained. Optionally, for top signals, Trex-QTL can be reapplied on *cis*-regressed data to variants that were previously pruned from the region to enable detailed characterization of candidate loci. The goal of this optional step is to characterize previously identified loci, rather than to discover new significant signals. **b.** Bottom: Genome-wide distribution of Trex-QTL scores in DGN across genomic positions (hg19), highlighting loci with strong global regulatory effects. Dark blue=significant lead variants (FDR<25%), light blue=significant variants in LD with a lead variant, gray=non-significant variant. Notable signals include loci near the previously identified *trans*-eQTL loci *ARHGEF3* (chr3:56849749) and *IKZF1* (chr7:50366637), which show elevated Trex-QTL scores indicative of widespread *trans* regulation. Top: Quantile–quantile (QQ) plots for the *ARHGEF3* and *IKZF1* loci comparing observed versus expected −log₁₀ p-values for *cis* (blue) and *trans* (orange) associations for each variant-gene pair, showing enrichment of *trans* signals detected by Trex-QTL.

### Genome-wide detection of trans-eQTLs in outbred rats

We applied Trex-QTL for *trans*-eQTL detection using 339 whole-brain RNA-seq samples from heterogeneous stock (HS) rats for which genotype data is also available^20^. HS rats are derived from eight genetically diverse inbred founder strains (**Fig. 3a**), and had been outbred for 77-85 generations at the time these samples were collected, resulting in genomes that are random mosaics of the eight founder haplotypes^21^. Since these rats have been bred in carefully controlled conditions and the samples were collected from healthy animals under treatment naive conditions, noise from environmental factors, which are prevalent in human datasets, is expected to be substantially reduced. Further, the breeding structure of this cohort has resulted in LD blocks that are larger than those observed in humans^20,21^, reducing the total number of tests needed to perform genome-wide association testing, albeit at the cost of fine-mapping precision. Overall, the relatively high rate of genetic diversity, high minor allele frequency (MAF) of polymorphic alleles, large LD block size, and lack of variation introduced by environmental factors leads to the HS rats dataset being better powered than human cohorts of comparable size for eQTL mapping^20^.

**Figure 3:**
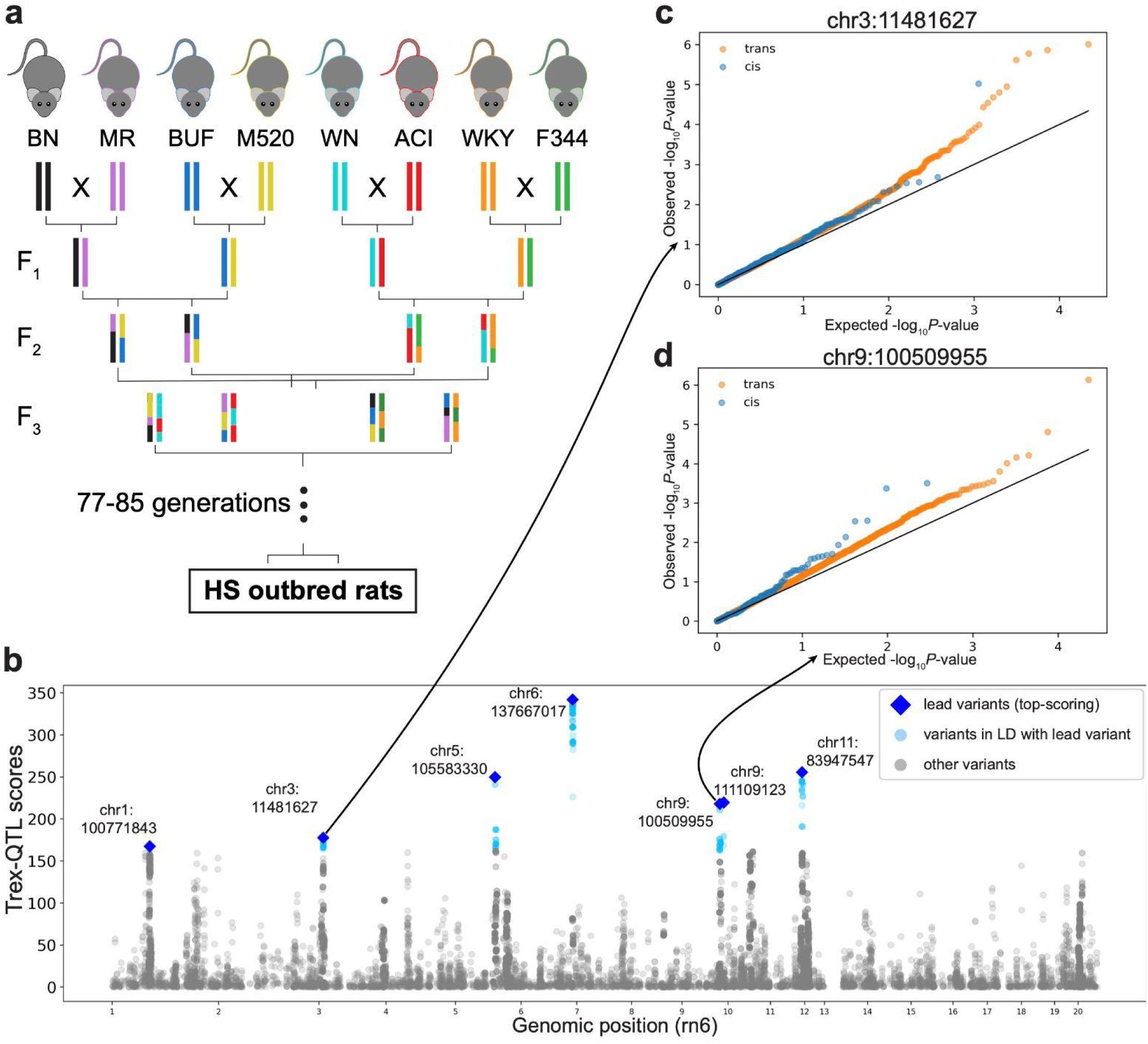
***Trans*-eQTL signals identified by Trex-QTL in HS rat brain samples. a.** A schematic depicting the establishment of the HS rat cohort. Outbred strains are derived from 8 inbred founder strains after breeding for 77-85 generations. **b.** Genome-wide distribution of *S* scores by genomic position (rn6), highlighting top-scoring loci. Dark blue=lead variants (top-scoring), light blue=variants in LD with a lead variant, gray=all other variants. **c-d.** Quantile-quantile (QQ) plots of association statistics across all genes for example top-scoring variants. Blue=-log10 *P*-values for genes in *cis* (same chromosome as the lead variant); Orange=-log10 *P*-values for genes in *trans* (different chromosome).

After data preprocessing, including removal of lowly expressed and poorly mapped genes, and exclusion of genes with highly heterogeneous expression patterns or strong global correlation structure (**Methods**), 11,745 protein-coding brain-expressed genes remained. In parallel, we obtained genotypes for approximately 6.6 million variants previously called using genotyping-by-sequencing^22^. To reduce the set of variants to those most likely to either influence protein function or alter expression levels of a potential *trans*-regulator, we restricted the analysis to variants within exons or within +/-3kb of the transcription start site (TSS) of a protein coding gene. We further removed variants with MAF < 0.1, >20% missing genotypes, or Hardy–Weinberg equilibrium P < 1×10⁻⁵ (**Methods**). We then applied LD pruning using PLINK to remove highly correlated nearby variants (within 100kb and r^2^>0.99), resulting in 11,000 stringently filtered variants for downstream analyses.

To improve power for *trans*-eQTL detection, similar to in the DGN analysis above, we first regressed out the effects of top *cis*-associated variants from gene expression levels (**Methods**; **Supplementary Fig. 6**). Even after this step, in the HS rat cohort we observed residual signal driven by *cis* variants, likely due to the larger LD blocks in this dataset compared to the DGN human data. Therefore to further mitigate effects that might be driven by residual *cis* signals, we excluded genes proximal to each variant when computing Trex-QTL statistics. Specifically, for each variant, we removed the *n* nearest genes such that the number of genes tested remained constant across variants (*n* = 303, corresponding to the smallest gene count on a single chromosome). All association analyses included sex, genotype principal components, and expression principal components as covariates.

Following these filtering steps, we next applied Trex-QTL to the *cis*-adjusted expression data (**Supplementary Table 1**). We identified top *trans*-eQTLs, defined as variants achieving the minimum attainable permutation-based *P*-value (P=1.82e-05, corresponding to <1/55,000), satisfying the hypothesis that target genes should show inflated summary statistics compared to a uniform distribution (estimated λ > 1), and representing lead variants after LD-based clumping (**Methods; Supplementary Table 2**). We removed from this set candidate *trans*-eQTLs driven by association with a single strong target gene that either falls on a sex chromosome or maps to multiple locations in the rat reference genome (rn6), since we hypothesize these may be driven by read mapping artifacts (removed *trans*-eQTL signals shown in **Supplementary Table 3**). After applying these filters, 7 strong candidate *trans*-eQTLs remained (**Fig. 3b**; **Table 1; Supplementary Fig. 7**). For each of these 7 candidate variants, inspection of pairwise gene-variant summary statistics showed the expected trend of enrichment for strong *P*-values for genes in *trans* (**Fig. 3c-d**; **Supplementary Fig. 8-9**).

**Table 1:** Top candidate *trans*-eQTLs identified in the HS rat cohort using Trex-QTL. Top, LD-independent *trans*-eQTLs identified following permutation-based significance testing, mixture model parameter filtering (λ > 1), and LD-based clumping (**Methods**). Variants are ordered by decreasing *S* score. Reported values include chromosomal position, *S* score, the predicted proportion of target genes (*t*), and the inferred mixture model parameter (*λ*), which reflects the estimated strength of departure of *P*-values for target genes compared to the expected uniform distribution. Loci shown represent high-confidence candidate *trans*-eQTLs remaining after filtering potential mapping artifacts and residual *cis*-driven signals that meet the minimum attainable *P*-value based on permutation testing (P<1.82e-05).

| Lead variant (rn6) | S | Predicted $t$ | Predicted $\lambda$ |
| --- | --- | --- | --- |
| 6:137667017 | 341.7 | 0.03 | 3.57 |
| 11:83947547 | 255.5 | 0.20 | 1.72 |
| 5:105583330 | 249.6 | 0.13 | 2.02 |
| 9:111109123 | 219.4 | 0.26 | 1.54 |
| 9:100509955 | 217.7 | 0.38 | 1.38 |
| 3:11481627 | 177.5 | 0.19 | 1.63 |
| 1:100771843 | 167.2 | 0.16 | 1.70 |

To evaluate whether these *trans*-eQTL loci may influence complex physiological and behavioral traits, we examined the top 7 lead variants to determine if other traits had shown genome-wide significant associations at the same or nearby variants in strong LD (R^2^>0.95; **Methods**). The signal with lead variant 6:137667017 showed 6 such associations including with the traits body length, weight of a specific leg muscle and two behavioral traits. Similarly, the signal with lead variant 5:105583330 showed 2 associations – one for BMI and one for a behavioral trait, and the lead variant 1:10077184 showed an association with kidney weight. These complex traits may be consequences of the widespread transcriptional changes induced by top *tran*s*-*eQTLs identified by Trex-QTLs.

### Top candidate trans-eQTLs implicate known gene regulatory pathways

To assess whether individual *trans*-eQTLs support a biological mechanism, we examined the top signals in more detail. The lead signal identified on Chromosome 6 spans an approximately 800 kb block of linked variants with strong *trans*-eQTL signals (**Fig. 4a-b**). Initially we inspected the functional consequences of protein-coding variants in this region, but did not identify any clear candidate variants (**Supplementary Table 4**). Next, we evaluated whether the lead variant in this region is a candidate *cis*-eQTL for genes that potentially mediate the *trans* signal. We found nominally significant (*P*<0.05) *cis*-eQTL effects for 11 expressed genes in the region (**Fig. 4c**). While the strongest effect is for *Nudt14* (*P=*6.59e-40), which is a Nudix hydrolase involved in UDP-glucose metabolism, we also observed a strong *cis*-eQTL for *Jag2* (*P=*2.22e-19), a critical ligand for the Notch pathway which is active in neural cells^23^. When *Jag2* binds to the Notch receptor, it triggers cleavage of the Notch Intracellular Domain (NICD), which translocates to the nucleus and interacts with a number of co-factors, including the DNA binding protein *Rbpj1* (also known as CSL)^24^ (**Fig. 4d**). We therefore hypothesized it has high potential to induce widespread *trans* effects.

**Figure 4:**
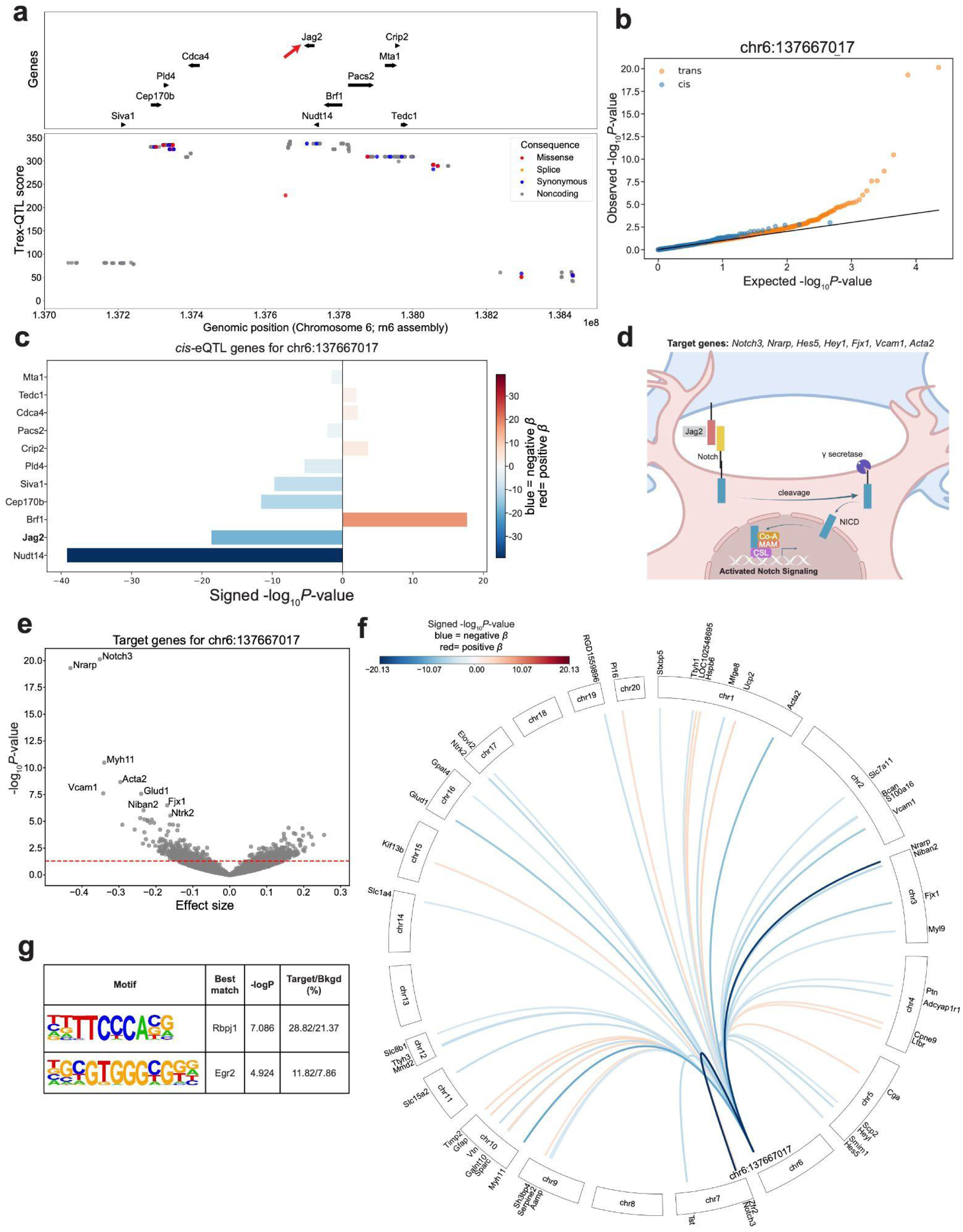
A *trans*-eQTL signal overlapping the Notch ligand *Jag2* is associated with Notch signalling pathway genes. **a.** *S* scores in the region surrounding the lead *trans*-eQTL variant. Color denotes the most severe functional consequences of each variant predicted by VEP. Dark blue=synomyous, orange=splice variant, red=missense, gray=noncoding. Only genes for which the *trans*-eQTL variant is also a significant *cis*-eQTL are shown. **b.** Quantile–quantile (QQ) plots for the lead variant chr6:137667017 (rn6) comparing observed versus expected −log₁₀ P-values for *cis* (blue; genes on the same chromosome) and *trans* (orange; genes on different chromosomes) associations, showing an enrichment of *trans* signals detected by Trex-QTL. **c.** All genes exhibiting nominally significant *cis*-eQTL associations within a ±500 kb window surrounding the lead variant (chr6:137,667,017). Bar length corresponds to the signed −log_10_(*P*), with negative values (blue) indicating decreased expression and positive values (red) indicating increased expression associated with the alternate allele. *Jag2* displays one of the strongest *cis* associations in the region and is highlighted in bold. **d.** A schematic showing the Notch signalling pathway, where the membrane bound *Jag2* ligand interacts with Notch, releasing NICD (the portion of the Notch receptor cleaved by γ-secretase) which translocates to the nucleus and activates gene expression with examples of target genes listed on the left. Red and blue cells indicate two interacting neurons. **e.** Volcano-plot summarizing associations between the lead *trans*-eQTL variant and each gene. The x-axis shows the effect size and the y-axis shows the association -log10 P-value. The red dashed horizontal line indicates nominal significance (P=0.05). **f.** Circos plot showing significant *trans*-eQTL associations for the lead variant at chr6:137667017. Each arc connects the variant to a target gene located on another chromosome. Arc color represents the signed association strength (signed −log10(P)); blue indicates that the alternate allele is associated with decreased gene expression (negative *β*), whereas red indicates increased gene expression (positive *β*). Only the top 50 target genes are shown for clarity. **g.** Motif analysis of promoters of 409 genes with negative effect sizes and Matrix-eQTL P<0.05 against 10,696 background genes. Columns from left to right show for each nominally enriched motif: sequence logo visualization, motif name, enrichment P-value, and % of target and background genes whose promoters contain the motif.

The top *trans*-eQTL target genes show a significant bias toward negative vs. positive effect sizes (409/746 genes with pairwise association *P*<0.05, binomial one-sided P=4.645e-03; **Fig. 4e-f**), indicating that the alternate allele at the Chr6 locus is associated with decreased expression of these genes. Top target genes contained many known Notch targets, including *Hes5*, *Hey1* and *Notch3*^25^ (Matrix-eQTL P-values: 5.794e-05, 7.416e-06, 7.376e-21; **Fig. 4e-f**). Motif enrichment analysis using HOMER^26^ of promoter regions (transcription start site -350bp/+50bp; **Methods**) of genes with pairwise Matrix-eQTL *P*<0.05 for which the *trans*-eQTL shows negative effect sizes identified the motif for *Rbpj1*, the central transcription factor in the Notch signaling pathway, as the top enriched known motif (**Fig. 4g**) which reached nominal significance (nominal -log P=7.086, FDR=0.37). Taken together, these findings are consistent with a model in which the *trans*-eQTL on Chromosome 6 reduces *Jag2* expression, which in turn decreases ligand-dependent activation of the Notch signaling pathway and reduces expression of downstream target genes. Intriguingly, the Notch signaling pathway has been implicated in two of the complex traits that were associated with this locus: muscle development^27^ and regulation of growth-plate chondrocyte differentiation and proliferation^28^, which is one of the determinants of longitudinal skeletal growth.

Another strong *trans*-eQTL locus was identified on Chromosome 5 overlapping a cluster of 8 interferon genes (**Fig. 5a; Supplementary Table 5**) which similarly showed evidence of enrichment for small P-values for *trans* associations with expression across the genome (**Fig. 5b**). Top target genes include genes *Parp14* and *C4a* whose expression is known to be regulated by interferon gamma^29,30^ as well as interferon regulatory factors *Irf7* and *Irf9* which are both activated by *Irf9* as part of the ISGF3 complex^31^ (**Fig. 5c-d**). Although the genes from the interferon cluster were not expressed in the brain samples analyzed (median TPM=0), we hypothesize that the variant(s) at this locus modulate interferon production in non-neuronal cells. In the brain, type I interferons are produced by glial cells and drive synapse loss^32^. We therefore propose that this locus acts through a cell non-autonomous fashion, where the allele(s) on Chromosome 5 impact expression of the interferon gene cluster in cells which are either not present, or represent a low proportion of the cellular population, in the brain samples used for this analysis, leading to changes in downstream signaling pathways (**Fig. 5e**).

**Figure 5:**
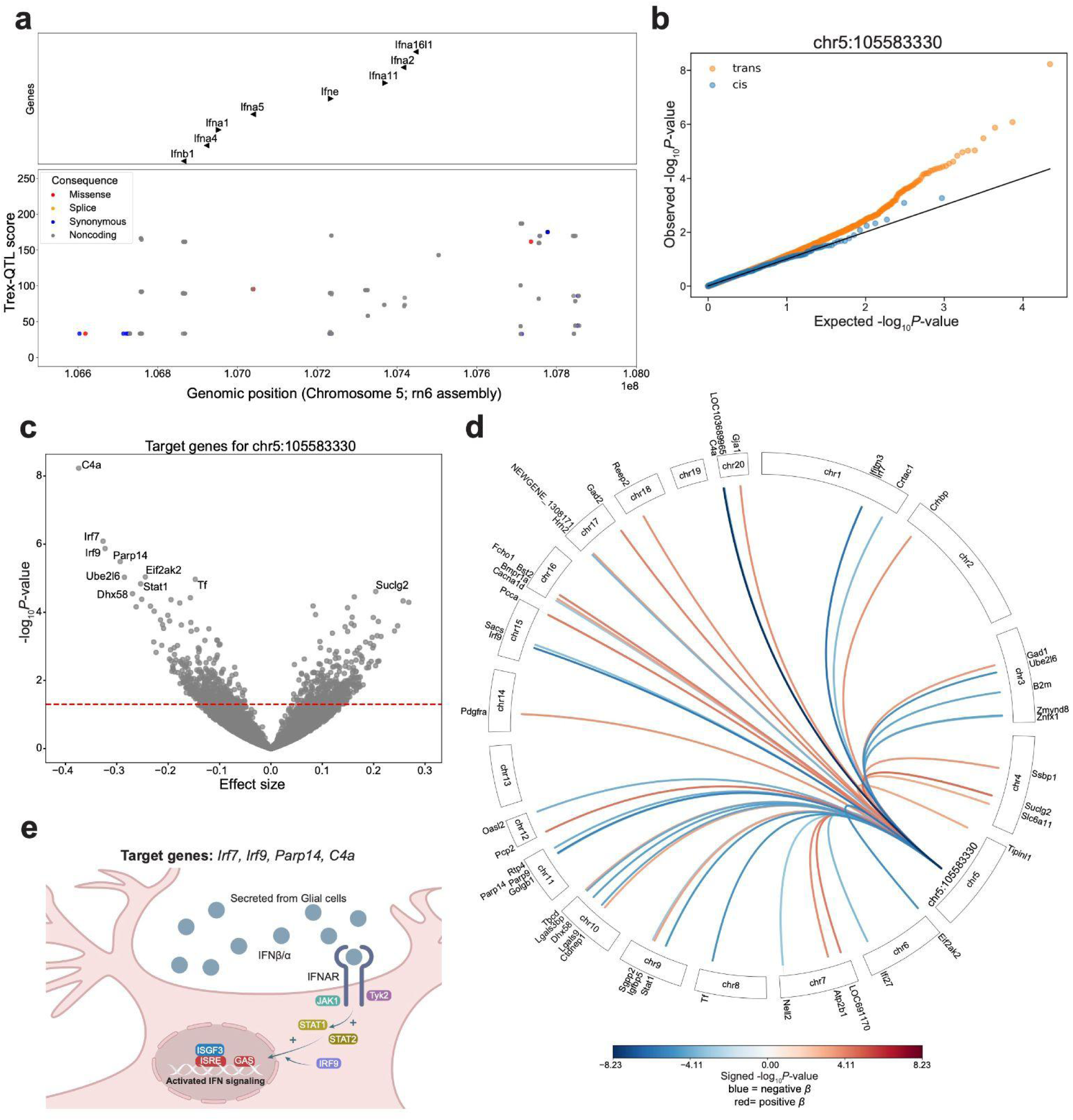
A putative *trans*-eQTL in the interferon cluster is associated with the Irf signalling pathway. **a.** *S* scores in the region surrounding the lead *trans*-eQTL variant. Color denotes the most severe functional consequences of each variant predicted by VEP. Dark blue=synonymous, orange=splice variant, red=missense, gray=noncoding. Only genes from the interferon cluster are shown. **b.** Quantile-quantile (QQ) plots for the lead variant Chr5:105583330 (rn6) comparing observed versus expected -log₁₀ p-values for *cis* (blue; genes on the same chromosome) and *trans* (orange; genes on different chromosomes) associations, showing an enrichment of *trans* signals detected by Trex-QTL. **c.** Volcano-plot summarizing associations between the lead *trans*-eQTL variant and each gene. The x-axis shows the effect size and the y-axis shows the association -log10 P-value. The red dashed horizontal line indicates nominal significance (P=0.05). **d.** Circos plot showing significant *trans*-eQTL associations for the lead variant at Chr5:105583330. Each arc connects the variant to a target gene located on another chromosome. Arc color represents the signed association strength (signed -log10(P)); blue indicates that the alternate allele is associated with decreased gene expression (negative *β*), whereas red indicates increased gene expression (positive *β*). Only the top 50 target genes are shown for clarity. **e.** Schematic showing the interferon pathway, where the INFB/a ligands bind to IFNAR on the neurons, inducing JAK/STAT signaling and Irf mediated gene expression changes, with a few examples of target genes listed on the left.

## Discussion

*Trans*-eQTLs have been notoriously difficult to detect due to insufficient statistical power with current sample sizes, relatively weaker effects compared to *cis*-eQTLs, and a lack of comparable tissues and cohorts for replication. We developed Trex-QTL, a novel *trans*-eQTL detection method that jointly models variant-gene summary statistics across all genes to identify variants with global impacts on expression of genes in *trans*. Compared to related methods, Trex-QTL employs a flexible model that can capture variants impacting either a few or thousands of target genes in *trans* and has reduced input requirements (summary statistics alone, or optionally individual-level data, rather than predefined gene modules). The novel, biologically focused, approach behind Trex-QTL is to model the effect of each *trans*-eQTL as a mixture of two distributions: genes that are targeted and ones that are not targeted by the variant. Specifically, Trex-QTL employs a mixture model of association statistics that simultaneously estimates the number of target genes and relative strength of effect of each variant while improving detection power across a range of scenarios. We additionally developed a simulation framework that enables analysis of the impact of various properties such as the number of target genes and effect size on *trans*-eQTL detection power. The framework also enables the inclusion of the effects of gene-gene correlation and covariates which we found to be critical for recapitulating patterns observed in real data. We used our simulation framework to benchmark Trex-QTL against another tool, CPMA, and observed improved power to detect *trans*-eQTLs with low number of targets or small effect sizes.

We demonstrated the utility of Trex-QTL by applying it to two real datasets. First, we analyzed whole blood expression data from the DGN cohort. Notably, we focused on this dataset since the majority of replicable *trans*-eQTLs in humans have so far been identified using whole blood or blood-related datasets^4,33^. While several pairwise *trans* variant-gene candidates have been identified in the Genotype Tissue Expression project (GTEx), most do not replicate across tissues^1^ and have not been observed in other datasets. Application of Trex-QTL to the DGN cohort demonstrated that the method is capable of detecting established *trans*-eQTLs in human populations. Top signals identified by Trex-QTL correspond to loci near *ARHGEF3* and *IKZF1*, two *trans*-eQTL hubs that have been independently identified using multiple approaches and replicated across blood and platelet datasets^4,11,18,19^. Replication of these established loci demonstrates that Trex-QTL is sensitive to biologically meaningful *trans*-regulatory effects.

While the DGN analysis primarily served as a replication of known *trans*-eQTLs, application of Trex-QTL to the heterogeneous stock (HS) rat cohort enabled discovery of novel *trans*-eQTLs and highlighted both the opportunities and challenges of *trans*-eQTL detection in model organisms. HS rats are derived from eight founder strains and have been maintained as an outbred population for dozens of generations, resulting in high genetic diversity, high minor allele frequencies, and genomes composed of founder haplotype mosaics (**Fig. 3a**). Combined with controlled environmental conditions, these features provide increased power for eQTL discovery relative to similarly sized human cohorts. At the same time, the genetic architecture of HS rats presents several analytical challenges. Extensive LD reduces the multiple testing burden and improves power to detect *trans*-eQTLs, but limits fine mapping resolution. As a result, the causal variants and genes underlying many *trans*-eQTL signals remain difficult to distinguish from nearby candidates. To mitigate this issue, we restricted analyses to variants with a high chance of functional impact, including variants within exons and variants proximal to transcription start sites. Future studies incorporating founder haplotype information, larger cohorts, or complementary functional datasets may help further refine causal mechanisms.

Overall, we identified 7 candidate *trans*-eQTL loci in the HS rats with global effects on gene expression. The two Trex-QTL signals that we highlighted provide mechanistic support through known biology and both also showed genome-wide significant associations with complex traits that have been measured in other HS rats. For the top Chromosome 6 locus, the *cis*-regulation of *Jag2*, consistent direction and significant enrichment of known downstream Notch target genes, and enrichment of the motif for *Rbpj1* in target gene promoters strongly implicates Notch signaling as the primary pathway driving this signal. RBPJ1 is the obligate transcription factor of canonical Notch signaling^34^, and Notch-Hes signaling plays essential roles in neural stem cell maintenance and fate specification^35^. Taken together, these data support a model in which the variant reduces *Jag2* expression, impacting ligand-dependent Notch activation in neural cells.

Interestingly, the candidate *trans*-eQTL in Chromosome 5 potentially illustrates a cell non-autonomous trans-regulation, where downstream interferon-stimulated genes (e.g., *Irf7*, *Irf9* and *Parp14*) are impacted despite the interferon gene cluster showing no expression in the bulk brain tissue used for in our study. We propose that the variant modulates interferon production on other cells, likely glial, and the resulting signaling drives expression of interferon-responsive genes in neurons, consistent with the established role of type I interferons in synapse loss^36^. This cell non-autonomous mechanism can likely explain other *trans*-eQTL associations in primary tissue data, and motivates single cell eQTL approaches to resolve the cell type specific origins of such effects.

The Trex-QTL method faces several limitations. First, while it enables estimation of the number of target genes, it does not allow the inference of specific target genes. Ultimately, larger datasets are needed to have sufficient power to implicate specific variant-gene associations. Second, it only accounts for the impact of a single variant on each gene. While our preprocessing step to regress out the impact of *cis* variants on gene expression somewhat mitigates this for *cis* effects, it does not allow for the possibility of multiple separate *trans* effects from different variants on the same gene. In future work it may be possible to apply elastic net or related approaches to regress out *trans* effects from other chromosomes prior to applying Trex-QTL. Finally, Trex-QTL itself cannot distinguish different technical vs. biological factors driving *trans* associations. While we regress out top expression PCs to mitigate the impact of technical factors, our simulations suggest this commonly used correction may be regressing out true *trans* signals. Further, we cannot rule out that individual *trans*-eQTL signals may be driven by differences in cell type proportion (e.g. vascular cells vs. neurons) across individuals. In future work we could apply methods such as CIBERSORT^37^ and explicitly model such proportions across samples. An additional future direction is to experimentally validate global *trans* effects e.g. using genome editing to introduce specific variants or knock out the candidate *trans* regulator.

Overall, Trex-QTL represents a powerful method for detecting *trans* regulators with global effects on expression or potentially other molecular phenotypes. Application to real datasets demonstrates the presence of such *trans*-eQTL hubs in both humans and model organisms and suggests they may represent a substantial source of phenotypic variation contributing to complex traits.

## Methods

### Overview of the Trex-QTL framework

Trex-QTL is a method to detect genetic variants with global effects on molecular phenotypes of interest. It takes as input summary statistics based on pairwise variant-phenotype association tests and outputs a scored list of candidate *trans*-QTLs. For each variant, it models summary statistics (-log *P*-values) for each pairwise variant-by-phenotype association test as a sample from a mixture distribution of null and non-null effects. It outputs the estimated percentage (*t*) of phenotypes (e.g. genes) that are targets as well as a *P*-value testing the null hypothesis that the variant has no effect on any phenotype. Notably, while we focus below on variant effects on gene expression (eQTLs), Trex-QTL can theoretically be used to detect global *trans* effects on other molecular phenotypes (e.g. metabolites).

We describe below preprocessing steps to obtain summary statistics followed by the main algorithmic steps of Trex-QTL. Alternatively, if summary statistics are already available users may skip the preprocessing steps and proceed directly to Trex-QTL.

### Preprocessing: Pairwise association testing

We first obtain association statistics (*P*-values) for each variant-phenotype pair, including both *cis* and *trans* effects. In practice, association statistics are computed by Matrix-eQTL^38^ assuming the model for each pair below.

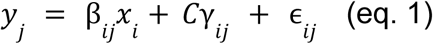

Where:

- *yj* is an *n* × 1 column vector of phenotype (e.g. expression) values for phenotype *j* in each of the *n* samples. Phenotype values are quantile normalized to follow a standard normal distribution.
- *xi* is an *n* × 1 column vector of genotypes for variant *i*. Genotypes are encoded as 0, 1, or 2, according to the count of the minor allele. Currently analysis is restricted to bi-allelic variants.
- β*ij* is a scalar equal to the effect size of variant *i* on gene *j*.
- *C* is an *n* × *c* matrix of covariates, where *c* is the number of covariates. Covariates often include sex, age, or technical sources of variation as measured by PEER factors^6^.
- γ*ij* is a *c* × 1 column vector of the effect sizes giving the effect of each of the *c* covariates on gene *j*.
- ɛ*ij* is an *n* × 1 column vector of error terms and represents variation in *yj* not explained by genotypes at variant *i* or covariates. Errors are assumed to be homoscedastic and independent and identically distributed, sampled from a normal distribution with zero mean.

#### Preprocessing: Removal of potential cis signals

To ensure we are detecting signals driven by *trans* (rather than *cis*) effects, we regress out potential *cis* effects from the molecular phenotype dataset. This step is optional for running Trex-QTL, however we have found that removing the effects of *cis*-eQTL can improve detection of *trans* signals in real data.

We implemented a stepwise conditional regression procedure to regress out significant *cis*-acting variants from each gene’s expression profile. For each gene, a user-specified *cis* window (*WindowSize*) centered on the gene’s midpoint was used to define candidate *cis*-variants. The optimal size of the window depends on the LD structure of the target dataset. For the rat dataset below, with large LD blocks, we initially set *WindowSize to* ±5 Mb. For the DGN human dataset, we set *WindowSize to* ±1 Mb. A set of candidate *cis*-variants for each gene was defined as those on the same chromosome and within the specified window.

To control for potential confounders, covariates such as sex, genotype PCs, and expression PCs are first regressed out of the expression data using ordinary least squares (OLS). The resulting residuals are then subject to an iterative conditional regression process to remove additive effects of cis-variants.

Specifically, for each gene:

1. Each candidate *cis*-variant is tested for association with the gene of interest using the covariate adjusted gene expression residuals using OLS.
2. The variant with the smallest *P*-value is identified.
3. If the *P*-value of the variant is less than 0.05, the variant is classified as a *cis*-eQTL and its effect is regressed out of the expression residuals for the gene.
4. The updated expression residuals are used for the next iteration.

This process is repeated up to a maximum of 10 iterations for each gene or until no additional variants meet the *P*-value threshold. For the HS rat dataset, the resulting residuals are used to perform pairwise association tests between variants and genes to identify genes exhibiting residual *cis* associations on the same chromosome that were not fully removed by the conditional regression procedure. The *WindowSize* is enlarged to ±25 Mb for these genes and the process to obtain expression residuals is repeated as above.

In the rat dataset, as an additional safeguard against residual *cis* contamination, genes proximal to each tested variant were excluded from Trex-QTL score computation below. To ensure that the number of genes evaluated was identical for all variants, we removed the nearest *n* genes surrounding each variant, where *n* was set to the smallest number of genes passing all filters present on any chromosome (n = 303, corresponding to Chromosome 11 in the HS rat dataset). This procedure minimizes the contribution of local *cis* effects while maintaining a consistent gene set size across variants, which enables using the same null distribution of *S* statistics across all chromosomes.

### Trex-QTL Step 1: Mixture model analysis

Under the null hypothesis that a variant is not associated with any of the molecular phenotypes, *P*-values of association statistics for variant-by-phenotype effects are expected to be uniformly distributed. Equivalently, under the null hypothesis, -log *P*-values are exponentially distributed with **λ**=1. However, if a variant is associated with a large number of phenotypes, this distribution will depart from the null, with **λ**>1.

A related method, CPMA^9^, models association statistics using a single distribution and detects variants for which **λ**>1. This model assumes all molecular phenotypes measured are targets of the variant of interest, whereas in practice a single *trans*-QTL likely only targets a subset of the phenotypes (e.g. expressed genes^39^). Instead, Trex-QTL models regression association statistics for each variant (-log *P*-values, denoted as *a* below) as a mixture of two distinct distributions (**Fig. 1a**, above) corresponding to target and non-target phenotypes:

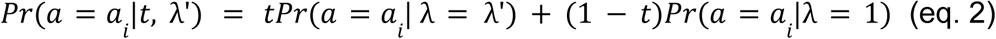

Where: *ai* is the i^th^ association statistic, *t* is the proportion of all phenotypes that are targets, 1/λ’ is the mean association statistic for target phenotypes, and *Pr*(*a* = *ai* | λ) = λ*e*^−λa^*^i^*. If the variant is not a *trans*-QTL, *t*=0, and the above model is equivalent to the null model in CPMA. By modeling effects as a mixture of null and non-null effects, Trex-QTL has sensitivity to detect a broader range of *trans*-QTLs, especially those affecting only a modest percentage of all phenotypes measured (supported by our simulation results in **Fig. 1**). Further, a key property of our model is that it can estimate the proportion of target genes (*t*) of a *trans*-QTL, a feature not enabled by alternative frameworks including CPMA.

Trex-QTL fits the mixture model above to obtain maximum likelihood estimates λ and *t* for λ’ and *t* using the Broyden–Fletcher–Goldfarb–Shanno (BFGS) algorithm^40^ based on the likelihood function:

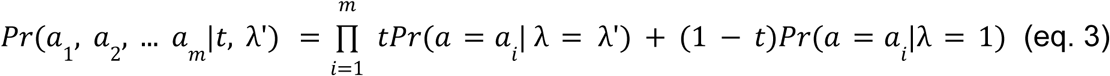

It then obtains a test statistic S using a likelihood ratio test to compare the best fit model to the null model with λ = 1:

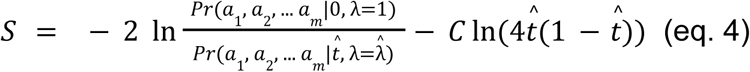

The second component of *S* above is a penalty term that shifts the mixture proportion *t* away from the boundary space of the parameter space (*t=0, t=1*) where standard likelihood ratio theory is not applicable. This penalty follows the modified likelihood framework proposed for finite mixture models by Chen and Kalbfleisch^41^. For Trex-QTL we set *C*=1 and found that the method to be robust to the choice of *C*.

### Trex-QTL Step 2: Significance testing

To obtain a significance value for *S*, we derive an empirical null distribution by simulating test statistics to recapitulate observed gene-gene correlations. To achieve this, we shuffled the sample labels in the genotype dataset, so they no longer correspond to those in the expression or covariate datasets. In this shuffled dataset, true *trans*-eQTL signals should be eliminated but gene-gene correlations are preserved. This shuffled genotype dataset is used to perform variant gene association testing and obtain Trex-QTL test statistics using the same steps as above. This process is repeated *x* times on the entire dataset to obtain a sufficiently large number of null values (*xq*) where *q* is the number of variants in the dataset. The resulting Trex-QTL test statistics are used as empirical null to obtain *P*-values on observed *S* values. A limitation of this approach is that the minimum possible *P*-value that can be obtained is 1/(*qx*). While performing more shuffling rounds can be used to achieve more precise *P*-values, this process is compute intensive. In practice we performed a total of 5 simulation rounds enabling obtaining *P*-values down to <1/55,000 for rats and 1 simulation round for *P*-values down to <1/1755712 for DGN.

### Simulation framework

To evaluate Trex-QTL, we developed a simulation framework to test various *trans*-QTL detection methods. The simulation framework follows a multivariate version of the regression model (eq. 1) described above considering the effect of a single *trans*-eQTL on *m* genes with *n* samples: *Y = X^T^β + ε*, where *X* is a length *n* vector of genotypes for the simulated variant, *β* is a length *m* vector of effect sizes of the variant on each gene, and *ε* is an *n*×*m* matrix of noise terms. Each row of *ε* is drawn from a multivariate normal *N*(0, *Cov*), with *Cov* set to the *m*×*m* identity matrix under default settings. The output of each simulation is an *n*×*m* matrix Y of expression values for each gene in each sample.

Unless otherwise specified, we set *m*=15,000 and *n*=500 to match the number of expressed protein coding genes and sample sizes of typical eQTL datasets. We vary the number of target genes and the effect size *β*. For non-target genes, *β* is set to 0. *β* is set to a constant non-zero value for all target genes. For each simulation, we additionally simulate data for 99 null variants with *β*=0 for all genes. We varied the number of target genes to range from 5-15,000 (ranging from 0.03% to 100% of all genes). Effect sizes varied from 0.01 (weak effects) to 1 (strong effects). Variant minor allele frequencies were set to 0.5. We also tested effect sizes and minor allele frequencies drawn from a normal distribution and found that doing so does not have a major impact on the comparison of *trans*-eQTLs detection methods. Thus, we chose a fixed effect size and minor allele frequencies to simply demonstrate the difference in power of *trans-*eQTL detection methods. For each simulated scenario of a specific *trans*-QTL setting, we performed 100 replicates.

The Trex-QTL simulation framework optionally allows inputting a gene-gene correlation matrix. If no matrix is provided all genes are assumed to be independent (*Cov*=*I,* where *I* gives the identity matrix). In our simulation experiments to account for the extensive correlation between expression of pairs of genes observed in real datasets^42^, we randomly simulate a gene correlation matrix *G*×*G* which is used in the simulation framework as ε ∼ *N*(0, *G*×*G*).

To assess the impact of covariates, we performed additional simulations including simulated effects of technical variation due to unknown sources, based on a technique previously published in the manuscript describing the PEER method^6^. We include 10 PEER factors in our simulations to demonstrate the effect technical covariates have on *trans*-eQTL detection methods. The model includes factor levels *l* and factor weights *w* for each simulated PEER factor:

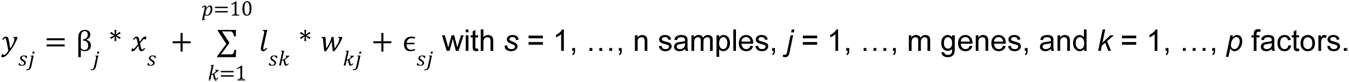

Factor levels *l*_sk_ for factor *k* are drawn from *N*(0, 0.6). Factor weights *w_kj_*of factor *k* for gene *j* were drawn from *N*(0, ^2^), where ^2^∼ 0.8(Γ(2.5, 0.6))^2^ which gives a heavy-tailed weight distribution.

### Simulation Power Analysis

To assess the power of *trans*-eQTL detection, we simulated expression datasets containing *trans*-eQTLs with varying effect sizes and numbers of target genes. The number of target genes ranged from 15 to 15,000, specifically: 15, 20, 30, 40, 60, 80, 100, 150, 200, 250, 300, 350, 400, 450, 500, 700, 1000, 2000, 3000, 4000, 5000, 6000, 7000, 8000, 9000, 10,000, 11,000, 12,000, 13,000, 14,000, and 15,000. The effect sizes were: 0.01, 0.02, 0.03, 0.04, 0.05, 0.1, 0.2, 0.3, 0.4, 0.5, and 1.

For each simulation, a single *trans*-eQTL was introduced using a unique combination of these parameters, and 100 datasets were generated for each parameter pair.

We then applied three *trans*-eQTL detection methods—Trex-QTL, CPMA, and Matrix-eQTLc—to each simulated dataset and evaluated their ability to detect the simulated signal. For Trex-QTL and CPMA, each simulation produced a single *P*-value summarizing evidence for a *trans*-eQTL, and power was defined as the proportion of simulations in which this *P*-value was below a specified significance threshold.

For Matrix-eQTL, which tests all variant–gene pairs, we first identified the set of true target genes based on the simulated effect sizes. Among these target genes, we extracted the corresponding variant-gene association *P*-values and used the minimum *P*-value per simulation as the detection statistic. Power was then defined as the proportion of simulations in which this minimum p-value passed the significance threshold.

Power was evaluated under multiple-testing-corrected significance thresholds expected in a genome-wide *trans*-eQTL study. For Matrix-eQTL, significance was defined using a Bonferroni correction based on 15,000 genes and 10,000 variants (α = 0.05/(15,000 × 10,000) = 3.33 × 10⁻¹⁰). For x-QTL and CPMA, which perform one test per variant rather than one test per variant–gene pair, we used a significance threshold of α = 1 × 10⁻⁴, corresponding to the approximate 10% false discovery rate observed in the HS rat dataset to provide a realistic threshold for genome-wide trans-eQTL detection.

For each parameter combination, we compared the empirical power across methods and recorded the method with the highest power to detect *trans*-eQTLs.

### Preprocessing expression and genotype data

#### Depression Genes and Networks (DGN) dataset

We used the preprocessed expression dataset available from the Depression Genes and Networks^15^ (DGN) study consisting of transcripts per million (TPM) values for each gene derived from whole-blood RNA-sequencing for 922 individuals of European ancestry. Genes with low expression (mean TPM < 0.1 across individuals) were excluded, resulting in 11,642 expressed genes retained for analysis. Expression values were log2-transformed and adjusted using all biological, technical, and hidden confounding factors provided with the DGN dataset. Specifically, we included all 75 covariates from the `Biological_and_hidden_factors.txt` and `Technical_factors.txt` files, comprising demographic variables (e.g., sex, age, body mass index, smoking status, medication use), genotype principal components (PC1–PC5), expression principal components (PC1–PC10), estimated blood cell composition, RNA-sequencing quality metrics, and other technical variables.

We used available genotype data from the DGN cohort and performed genotype imputation using Beagle v5.5^43^ with the 1000 Genomes Phase 3 reference panel (GRCh37/hg19). Standard quality control filters were applied to remove variants with Hardy–Weinberg equilibrium *P*-value < 1 × 10⁻⁶, MAF < 5%, or call rate < 95%.

#### Heterogeneous Stock (HS) rat dataset

We utilized available RNA-sequencing from the brain hemisphere of 339 HS rats^20^, aligned to the rat reference genome assembly rn6. Gene annotations were obtained from the Ensembl release 99 gene annotation (Rattus_norvegicus.Rnor_6.0.99.gtf.gz), available from the Ensembl FTP archive. The expression dataset consists of TPM values for 32,576 genes. Genes were filtered to remove those with no expression variability (variance = 0 or IQR = 0), extremely high variability (variance ≥ 50,000), low expression (median TPM ≤ 2 or maximum TPM ≤ 2), and non-protein-coding genes. We also removed genes located in segmental duplication regions. For this, we obtained segmental duplications annotated in the rn3 assembly from the UCSC Genome Browser^44^ and used the UCSC liftOver tool^45^ to convert coordinates to rn6. Gene coordinates from the Ensembl GTF annotation were then compared with the lifted-over segmental duplication intervals, and genes with any genomic overlap were removed. We additionally filtered genes with highly correlated expression profiles, and genes exhibiting excess heterozygosity as described below.

To detect genes with high heterozygosity, we calculated heterozygous genotype counts from polymorphic variants in the HS founder rats. For each gene, we performed a one-sided binomial test, treating each nucleotide in the gene as a trial and the observed number of heterozygous genotype calls as successes. Under the null hypothesis, the probability of success was equal to the genome-wide ratio of heterozygous genotype calls to total gene length across all analyzed genes.. *P*-values were Bonferroni-corrected, and genes with adjusted *P*-value < 0.05 were excluded. To identify highly correlated genes, we computed pairwise Spearman correlations across all genes. We then built a graph in which nodes represent genes and edges connect pairs of genes with absolute correlation>0.99. We retained a single representative gene per connected component. After filtering, the expression dataset contained 11,745 genes. The filtered expression matrix was then quantile normalized to standard normal distribution for each gene

Genotype calls initially consisted of 6,621,609 variants. Variant-level filtering removed low-frequency variants (minor allele frequency < 0.1), variants with >20% missing genotypes, variants failing Hardy-Weinberg equilibrium testing (P < 1×10⁻⁵), and multiallelic variants. We additionally excluded 10,144 variants located within segmental duplication regions.

To prioritize variants with potential regulatory or functional effects, we restricted analyses to variants located within exons or within ±3 kb of transcription start sites of protein-coding genes, removing 3,604,659 variants. We then applied LD pruning using PLINK v1.90b3.44^46^ (--indep-pairwise 100 10 0.99), removing an additional 162,666 variants. Finally, we excluded 211 variants lacking sufficient genotype diversity (fewer than three unique diploid genotypes with at least three samples each). The final dataset contained 11,000 variants for downstream analyses.

Expression values were adjusted for sex, the top five genotype PCs, and the top 20 expression PCs prior to regressing out *cis*-eQTL effects. Expression PCs were computed on the filtered expression data after normalization. Genotype PCs were obtained from the RatGTEx website (https://ratgtex.org/download/v1/).

### Genomic control

For the DGN dataset, we observed overall inflation in the distribution of unadjusted *P*-values of pairwise variant-gene pairs, suggesting residual confounding or unmodeled technical variation. To address this, we implemented a genomic control procedure^47^ based on the assumption that -log *P*-values follow an exponential distribution under the null hypothesis of no association. Specifically, we estimated the genomic control factor as the ratio of the observed median of -log(*P*) to its expected null median (log(2)) (equivalently, −log(0.5)). Each -log(*P*) value was then divided by this factor, and the adjusted *P*-values were obtained by exponentiation back to the *P*-value scale. This approach is equivalent to the classical χ²-based genomic control, but adapted for the exponential distribution of -log(*P*). This adjustment reduced inflation and improved calibration of the null distribution, thereby reducing the rate of false positives while retaining power to detect real *trans*-eQTL signals. We applied this correction only to the DGN dataset, as the HS rat dataset showed no evidence of inflation.

### Identifying top signals

For the DGN dataset, we first selected all variants that passed a 25% false discovery rate (FDR) using the Python module statsmodels.stats.multitest.multipletests^48^ using the Benjamini-Hochberg method^49^ with method option “fdr_bh”. threshold based on Trex-QTL results. To ensure comprehensive coverage of association signals, we then reintroduced variants that had been pruned out but were in LD with these significant variants. Trex-QTL analysis was repeated on this expanded set of variants, and those that again passed the 25% FDR threshold were retained. To identify the most representative variants for downstream analyses, LD-based clumping was performed using PLINK v1.90b3.44^46^ (--clump-kb 1000), ranking variants by Trex-QTL scores.

For the HS rat dataset, we applied a similar strategy with modifications to account for differences in genetic architecture and resolution. Because this cohort is characterized by large LD blocks with many nearby variants showing *P*-values at the minimum possible threshold, we instead selected variants with *P*-values equal to the smallest possible value, 1/(xq), where x is the number of permutations and q is the number of variants tested. In addition, only variants with estimated λ > 1^ were retained, as we expect that variants with λ ≤ 1 may correspond to technical or stochastic artifacts. The retained variants were then subjected to PLINK v1.90b3.44^46^ clumping (--clump-kb 5000) then ranked by Trex-QTL scores to define top independent hits. As an additional quality control step, we manually excluded loci whose signals were dominated by a single strongest associated target gene located on sex chromosomes or that mapped to multiple genomic locations, as determined by the Rat Genome Database (RGD) genome browser, since this can indicate mapping ambiguity issues or highly repetitive sequence content. Signals removed by these additional steps are listed in **Supplementary Table 3**. We additionally removed loci whose strongest target genes corresponded to ribosomal genes or pseudogenes, as these gene classes are particularly susceptible to read-mapping artifacts and ambiguous quantification. Notably, some pseudogenes remained annotated as *protein_coding* in the rn6 gene annotation used for this study and therefore were not removed by the initial protein-coding gene filter. To further reduce the possibility of residual *cis* effects, loci whose dominant target gene was on the same chromosome as the associated variant were also excluded.

### Evaluating associations with additional phenotypes in HS rats

We previously performed a phenome-wide association study (PheWAS) in the HS rat cohort. We identified lead Trex-QTL variants that are in LD (R2>0.95) with a genome-wide significant variant from PheWAS, defined as -log10(P)>5.6. We collapsed traits known to be highly phenotypically correlated (e.g. ’right kidney weight’ and ’left kidney weight’) such that those were only counted once.

### Interrogating top signals

To characterize the top *trans*-eQTL loci identified by Trex-QTL, we performed downstream analyses to define target genes, visualize association patterns, and assess potential regulatory mechanisms. For each candidate *trans*-eQTL variant, we considered genes for which nominal pairwise association *P*-values from Matrix-eQTL were < 0.05 as candidate target genes. Target gene sets were further stratified by direction of effect (positive or negative *β*). Only genes considered in the Trex-QTL computation (i.e. excluding the *n*=303 nearest genes to each variant) were considered as potential target genes. We identified candidate *cis*-associated genes (nominal *P*-value < 0.05) within ±500 kb around each variant. These genes were used to contextualize potential local regulatory effects at each locus.

Functional consequences of the variant set were annotated using Ensembl Variant Effect Predictor^50^ (VEP v97.3) with the Ensembl release 97 rat database for the Rnor_6.0 (rn6) reference genome assembly. Variants were classified according to their predicted consequences (e.g., missense, synonymous, splice-region, and noncoding variants), and these annotations were used for visualization and interpretation of candidate trans-eQTL loci.

### Motif enrichment analysis

To investigate potential regulatory mechanisms, we performed motif enrichment analysis on sets of upregulated, downregulated, and all target genes using HOMER^26^ (findMotifs.pl). Background genes were defined as all genes included in the expression dataset after filtering. Genes were supplied directly to HOMER, which performed motif enrichment analysis using its default promoter-based workflow, searching sequences surrounding transcription start sites (TSSs) in the rn6 reference genome. Enrichment of known transcription factor motifs was assessed relative to the background gene set using HOMER’s default parameters.

## Supporting information

Wu_etal_SuppMaterial

Wu_etal_SuppDataset1

Wu_etal_SuppTables

## Declarations

### Ethics and consent to participate declarations

Not applicable.

## Ethics declaration

Not applicable.

## Code availability

The Trex-QTL software package, including the simulation framework used in this study, is available at GitHub: https://github.com/cynthiaewu/trans-eQTL. The repository contains installation instructions and documentation required to run Trex-QTL. Analysis scripts and Jupyter notebooks used to reproduce the figures and analyses are available at https://github.com/cynthiaewu/TREX-manuscript.

## Availability of data and materials

Trex-QTL summary statistics for the DGN analysis are provided in **Supplementary Dataset 1.**

Summary statistics for HS rat analysis are provided in **Supplementary Table 1**.

DGN data were accessed through the NIMH Center for Collaborative Genomic Studies on Mental Disorders, under the ‘Depression Genes and Networks study’ (D. Levinson, PI).

## Acknowledgements

We thank Rahel Wachs for her help with the illustrations. Data for this publication was in part obtained from NIMH Repository & Genomics Resource, a centralized national biorepository for genetic studies of psychiatric disorders.

## Author contributions

C.W. implemented Trex-QTL, performed all analyses, and wrote the manuscript. A.B. helped with mathematical details of the Trex-QTL method and provided input on the analyses and manuscript. T.X. provided input on analyses and helped prepare the manuscript. E.M.M. helped supervise analyses and with biological interpretation of top signals, and helped write the manuscript. H.C. generated rats and collected brain samples.

F.T. processed brains for RNA extraction and generated RNAseq libraries and sequencing data. O.P. provided project management for the rat genotyping, tissue storage, and data organization. D.M. processed the rat RNA-seq dataset. A.A.P. helped supervise analyses, obtain funding, provide domain-specific expertise on HS rats, and with writing and editing the manuscript. A.G. co-supervised analyses, obtained funding, provided biological expertise for interpretation of signals, and wrote the manuscript. M.G. conceived the Trex-QTL method, co-supervised analyses, obtained funding, and wrote the manuscript.

