## Supplementary material for "Trex-QTL: A mixture-model for identification of genetic effects with global effects on molecular phenotypes": Wu_etal_SuppMaterial

### Supplementary Figures

#### Supplementary Fig. 1

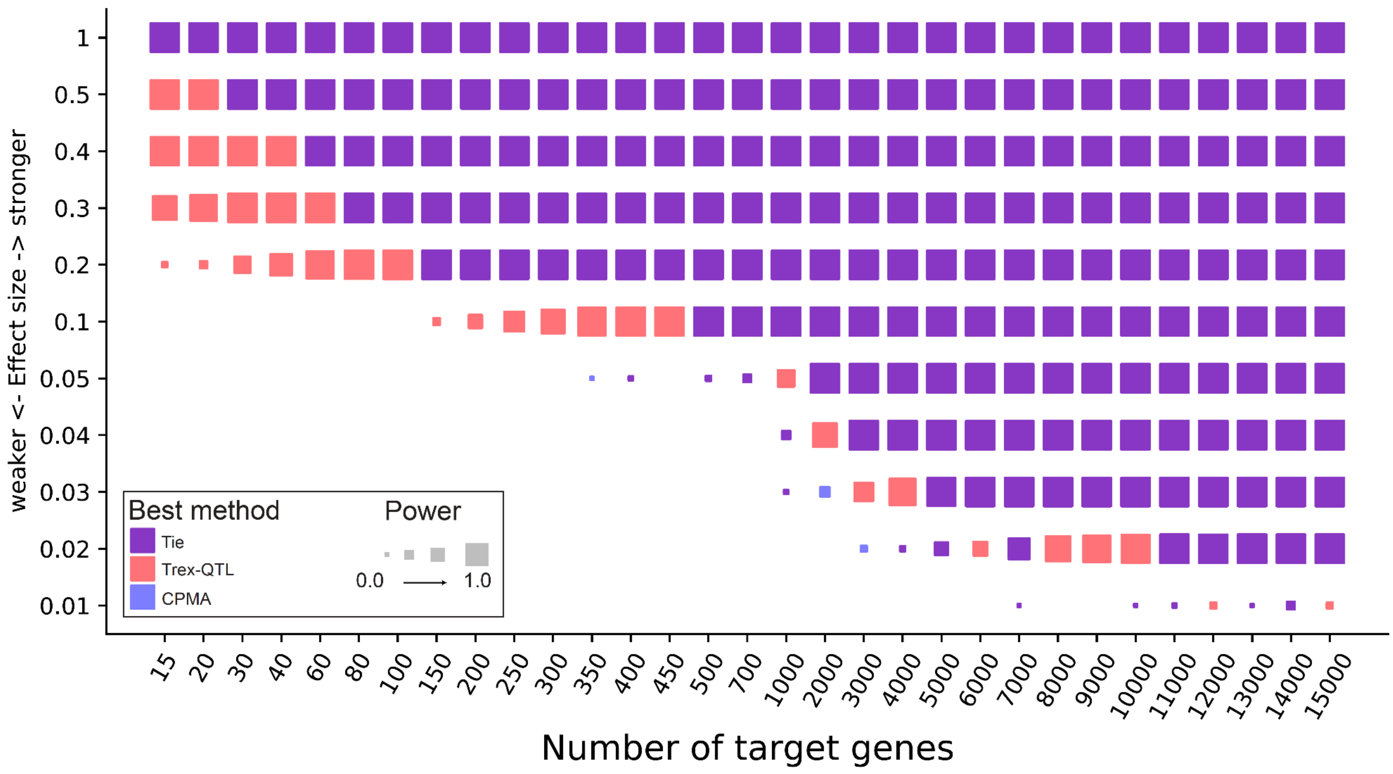


**Power comparison between Trex-QTL and CPMA.** Results from our simulation framework represented as a heatmap comparing Trex-QTL vs CPMA. The heatmap shows the *trans*-eQTL detection method with highest power for detecting *trans*-eQTLs with varying number of target genes and effect sizes. The color represents the method with highest power: blue=CPMA, red=Trex-QTL, purple=tie between Trex-QTL and CPMA. Both methods achieve high power for trans-eQTLs with large effect sizes and/or many target genes. However, Trex-QTL outperforms CPMA in scenarios with smaller effect sizes and/or fewer target genes, whereas CPMA shows modest advantages in a limited number of parameter settings.

#### Supplementary Fig. 2

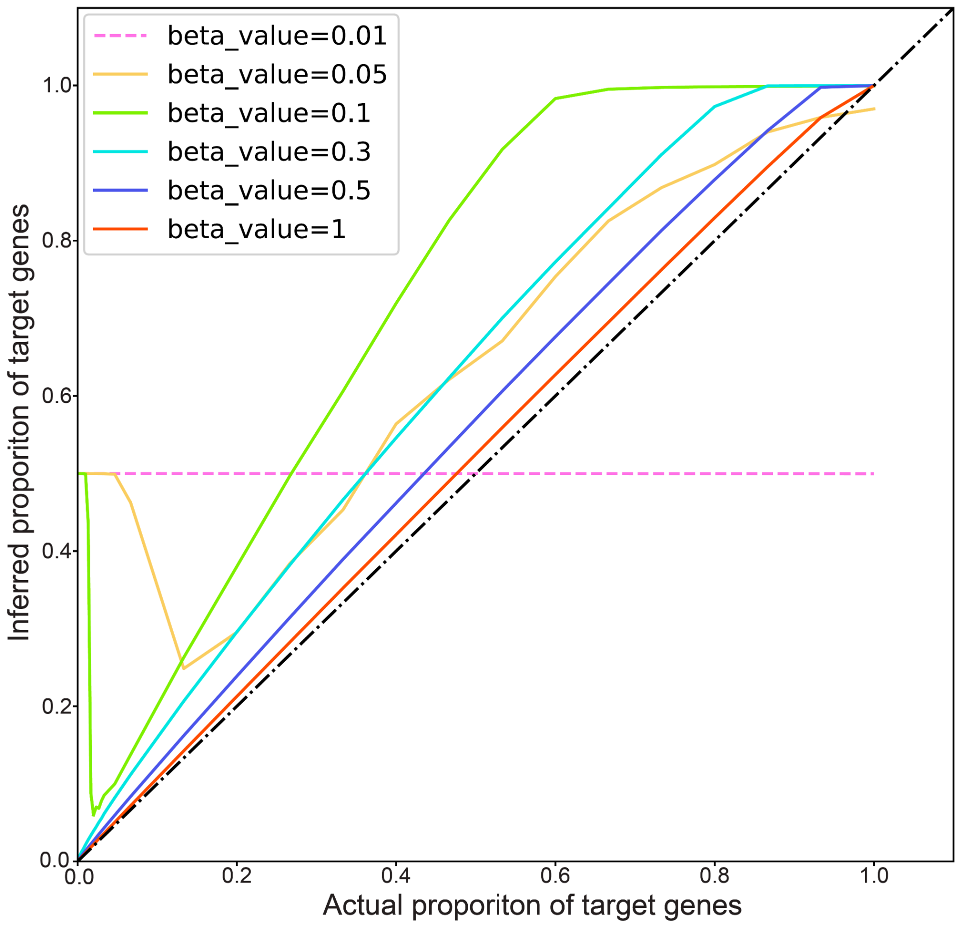


**Accuracy of Trex-QTL estimates of the proportion of target genes (*t*) in simulated datasets.** For each simulation, *trans*-eQTLs were generated with varying true proportions of target genes (*t*; x-axis) and effect sizes (β; colored lines). The y-axis shows the proportion of target genes inferred by Trex-QTL. The dashed diagonal indicates perfect agreement between the simulated and inferred values. Across moderate to large effect sizes (β ≥ 0.05), inferred *t* generally tracks the true proportion of target genes, although estimates tend to be inflated at larger values of *t*. For very weak effects (β = 0.01), Trex-QTL has limited power to accurately estimate *t*.

#### Supplementary Fig. 3

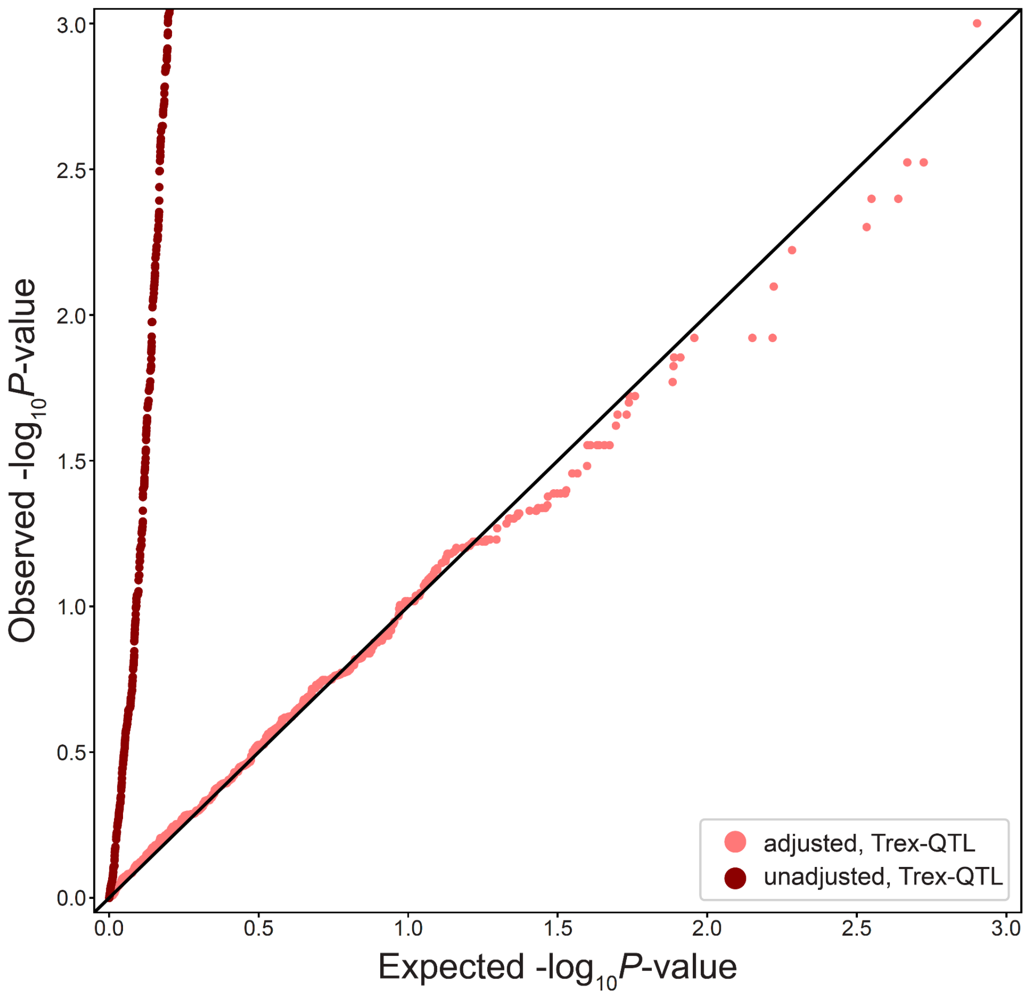


**Empirical null distributions correct p-value inflation caused by gene–gene correlation.** QQ plots of Trex-QTL p-values from 1,000 simulated null variants in a dataset with gene-gene correlation. P-values computed using a theoretical null distribution assuming independent gene-level tests (dark red) are strongly inflated, whereas p-values obtained from a permutation-based empirical null distribution (light red) are well calibrated and closely follow the expected distribution. Gene–gene correlation therefore necessitates empirical significance testing for accurate *trans*-eQTL detection.

#### Supplementary Fig. 4

**a**

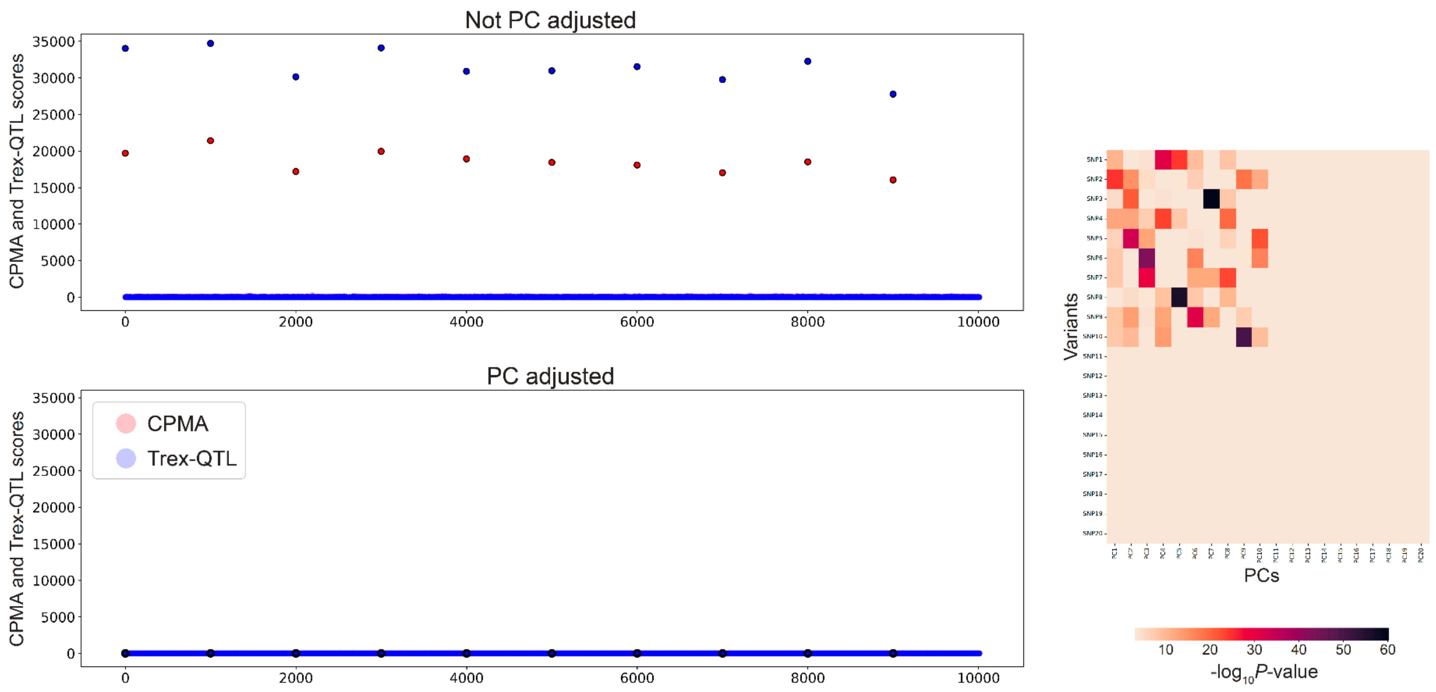


**b**

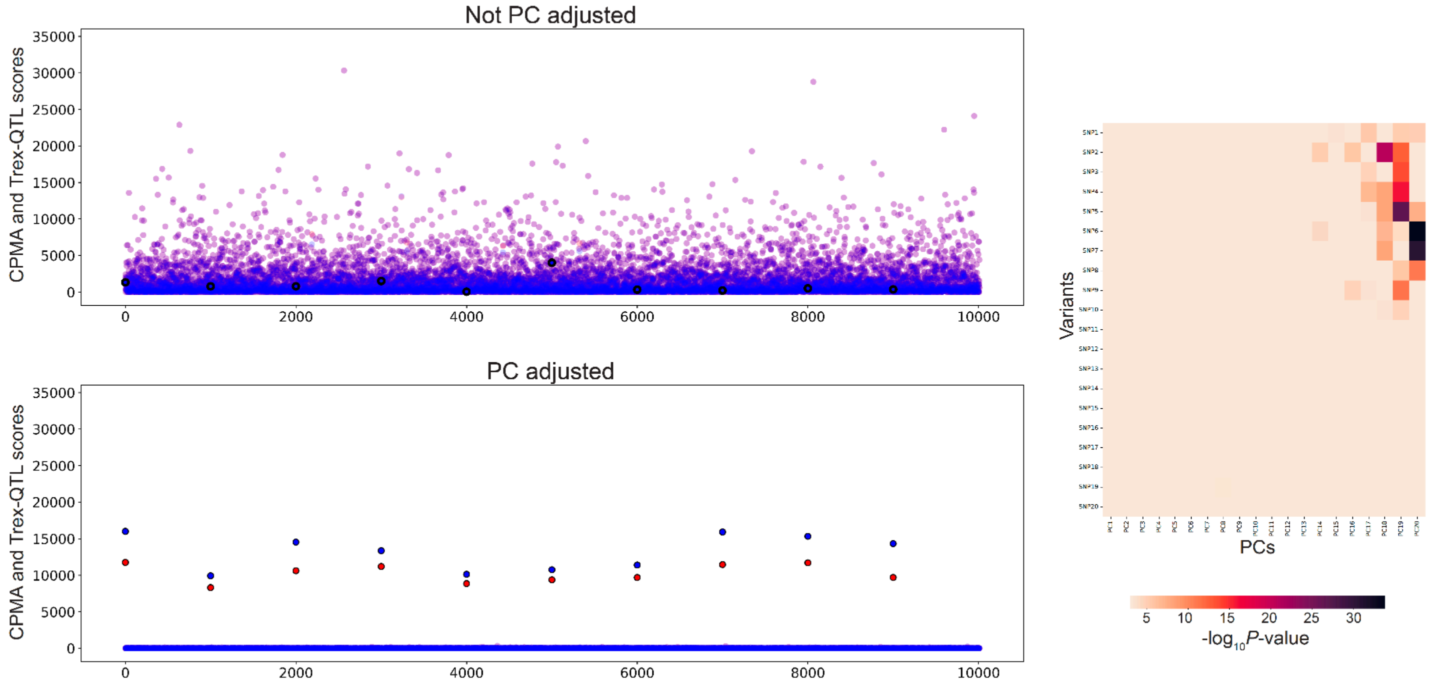


**Tradeoff of *trans*-eQTL analyses with or without PC adjustment.** The Manhattan plots (left) represent various simulation and PC adjustment scenarios. The x-axis corresponds to the 10,000 simulated SNPs. The

10 simulated *trans*-eQTLs (β=0.5 for 1,000 target genes out of 13,000 genes) are located at intervals of 1,000 starting from 0. The y-axis represents the CPMA and Trex-QTL scores of each variant. The heatmaps (right) plot the -log(p-value) of pairwise association testing of variants and PCs. The x-axis represents the first 20 PCs of the expression dataset. The y-axis corresponds to 20 variants. Variants 1-10 are the simulated *trans*-eQTLs and variants 11-20 are null. **a.** No PEER factors were simulated. *Left:* In the *trans*-eQTL analysis without PC adjustment, we can identify the 10 simulated *trans*-eQTLs. In the run with PC adjustment, we cannot identify the simulated *trans*-eQTLs. *Right:* The heatmap shows the top 10 PCs correspond to the 10 trans-eQTLs. PCs do not correspond to null variants. **b.** 20 PEER factors were simulated. *Left:* In the *trans*-eQTL analysis without PC adjustment, we cannot identify the simulated *trans*-eQTLs. In the run with PC adjustment, we can identify the 10 simulated *trans*-eQTLs. *Right:* The heatmap shows PCs 14-20 correspond to the 10 trans-eQTLs. PCs do not correspond to null variants.

#### Supplementary Fig. 5

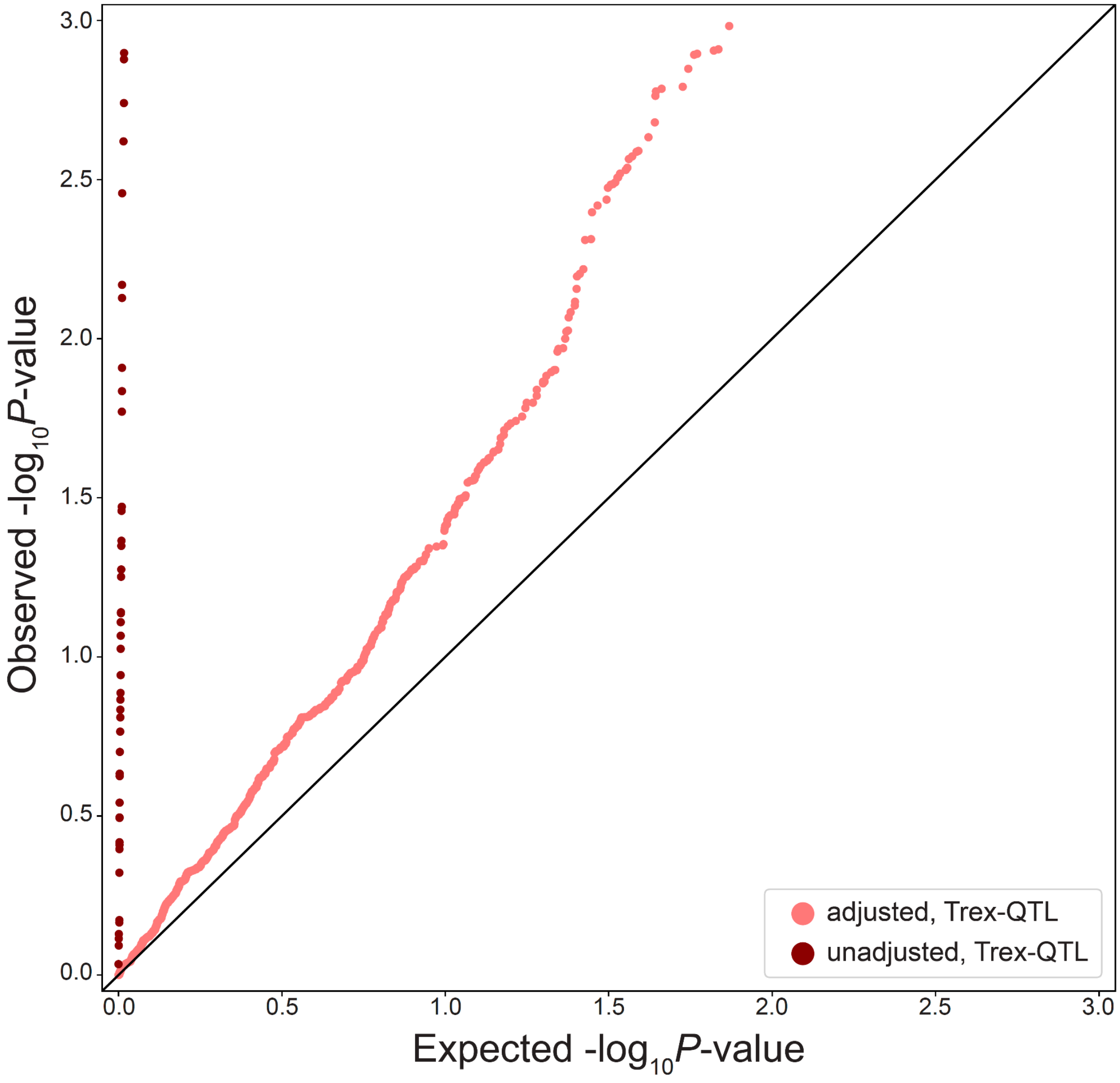


**QQ plot of Trex-QTL association statistics after correction for simulated technical variation.** QQ plots of observed versus expected −log10(P) values for Trex-QTL association statistics computed with (light red) or without (dark red) inclusion of the top 10 expression PCs as covariates in the presence of simulated 10 PEER factors in the expression dataset. Adjusting for expression PCs substantially reduces inflation of the test statistics, although moderate inflation remains, motivating the use of a permutation-based empirical null distribution.

#### Supplementary Fig. 6

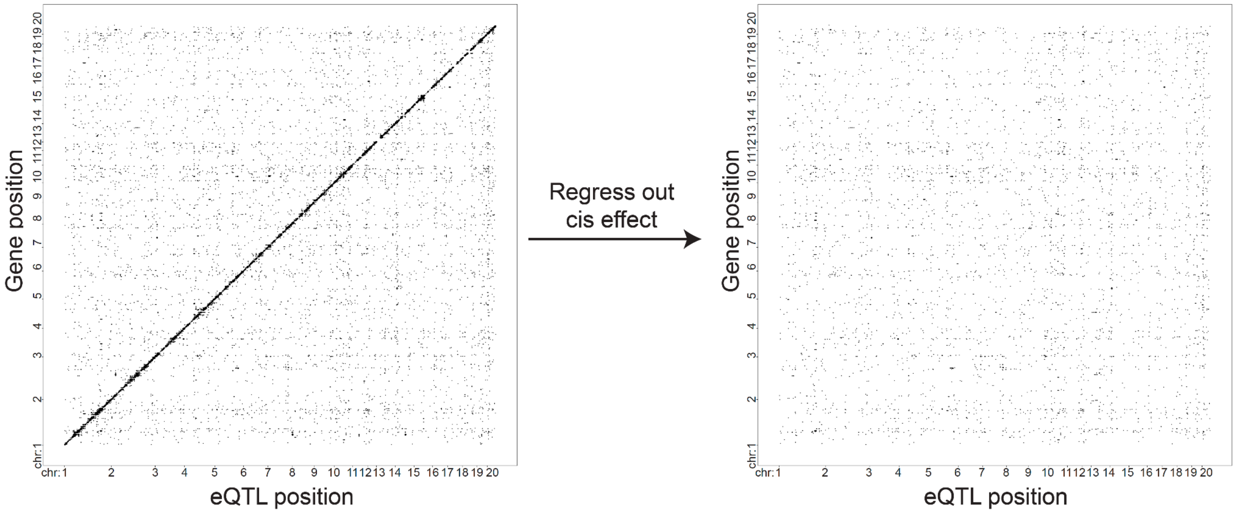


**Removal of cis-eQTL signals from the HS rat expression dataset.** Pairwise variant-gene association statistics were computed using Matrix-eQTL before (left) and after (right) application of the *cis*-regression procedure. The x-axis denotes variant genomic position and the y-axis denotes gene genomic position. Only associations with $P < 5 \times 10^{-5}$ are shown. Point size is proportional to $-\log(P)$, such that larger points indicate stronger associations. Prior to *cis*-regression, a strong diagonal pattern is observed, reflecting widespread *cis*-eQTL associations between nearby variants and genes. Following iterative regression of top *cis*-associated variants from gene expression levels (**Methods**), the diagonal signal is largely eliminated, indicating successful removal of *cis* effects while retaining distal *trans*-association signals.

### Supplementary Fig. 7

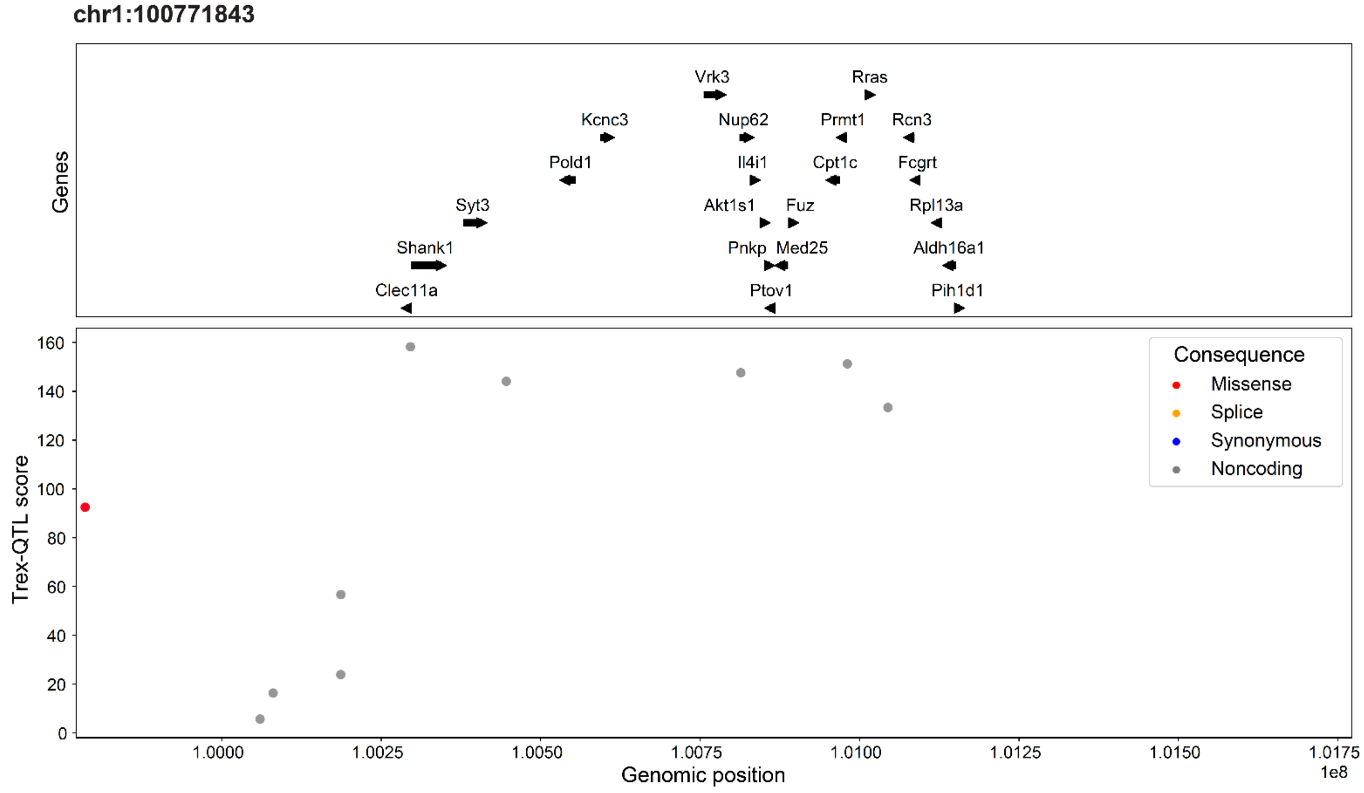


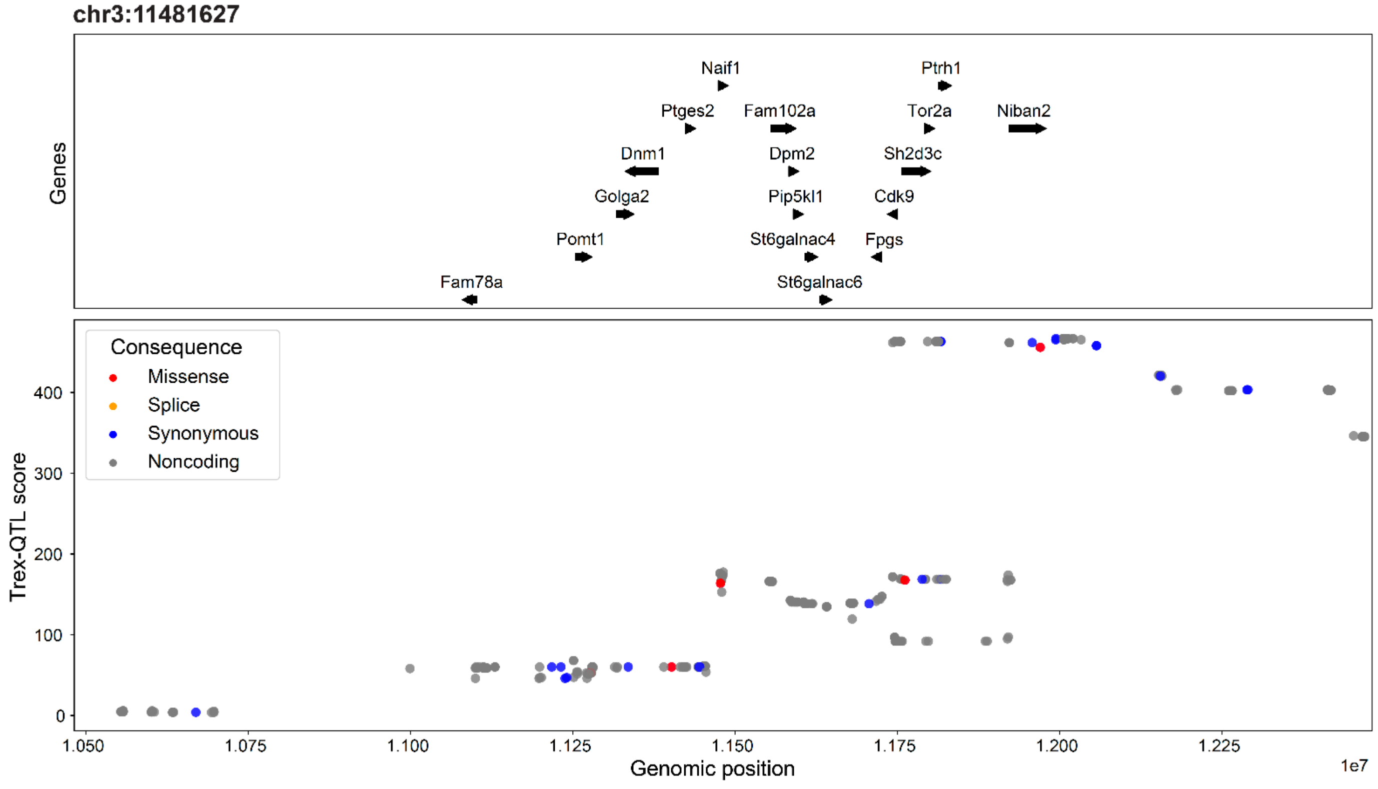

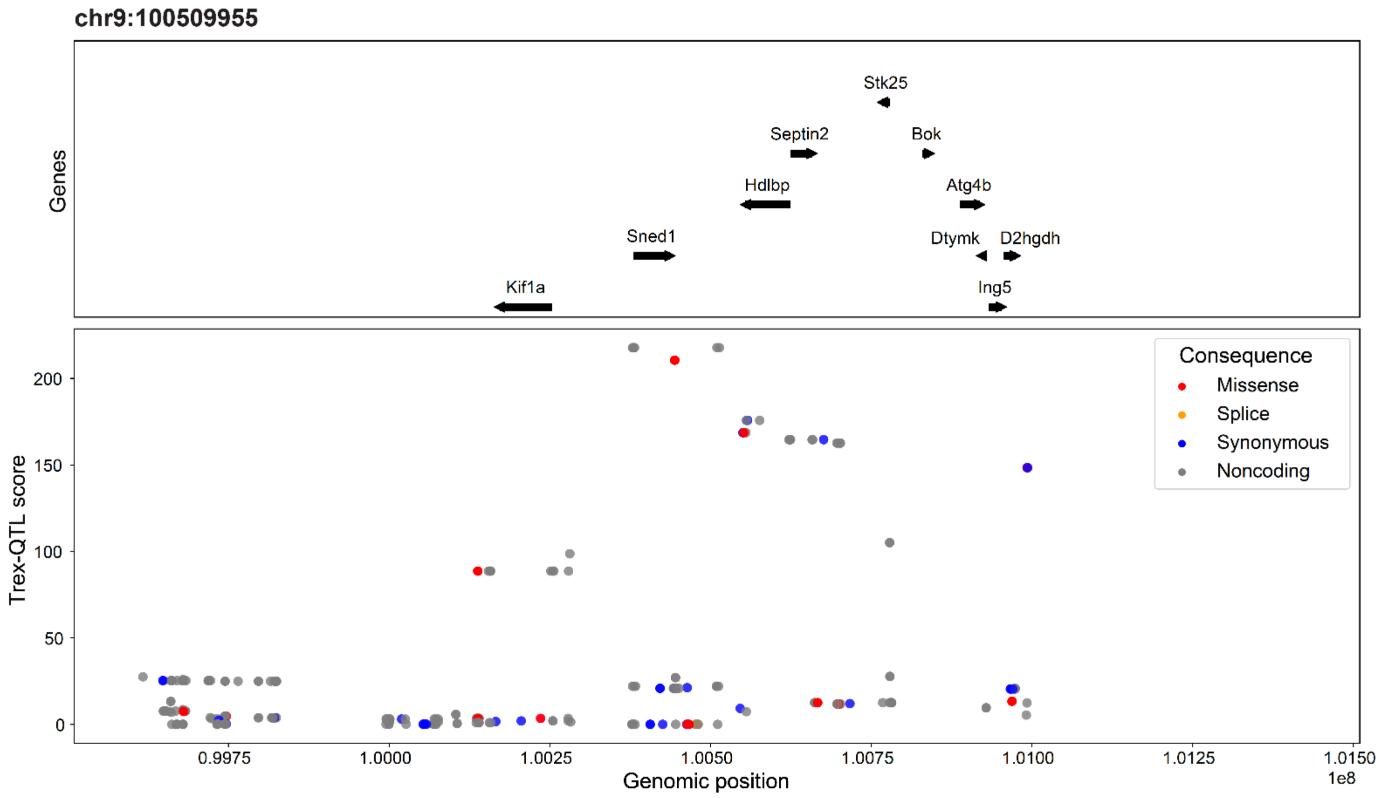
chr9:100509955
Genes
Stk25
Septin2
Bok
Hdlbp
Atg4b
Sned1
Dtymk
D2hgdh
Kif1a
Ing5
Trex-QTL score
Consequence
Missense
Splice
Synonymous
Noncoding
0
50
100
150
200
0.9975
1.0000
1.0025
1.0050
1.0075
1.0100
1.0125
1.0150
1e8
Genomic position

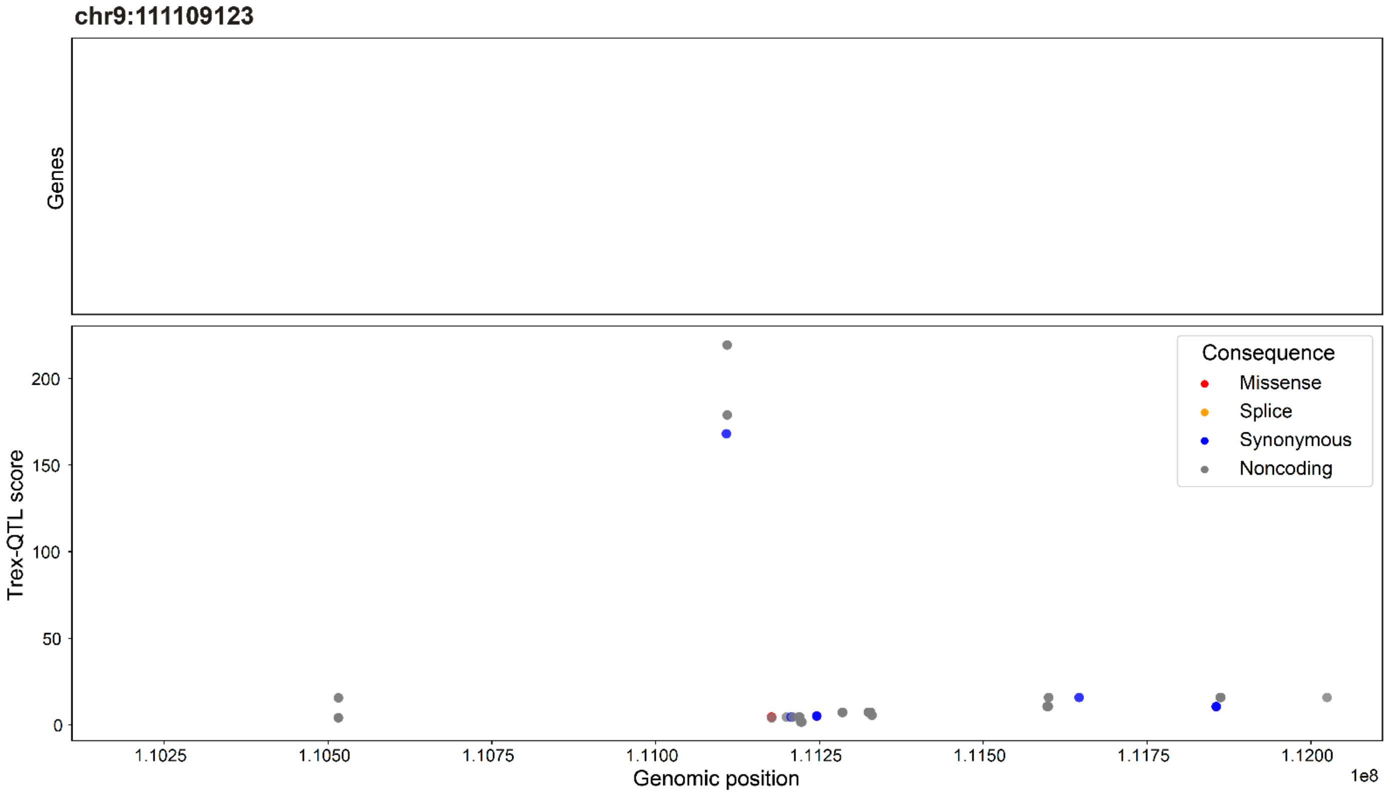
chr9:111109123
Genes
Trex-QTL score
Consequence
Missense
Splice
Synonymous
Noncoding
0
50
100
150
200
1.1025
1.1050
1.1075
1.1100
1.1125
1.1150
1.1175
1.1200
1e8
Genomic position

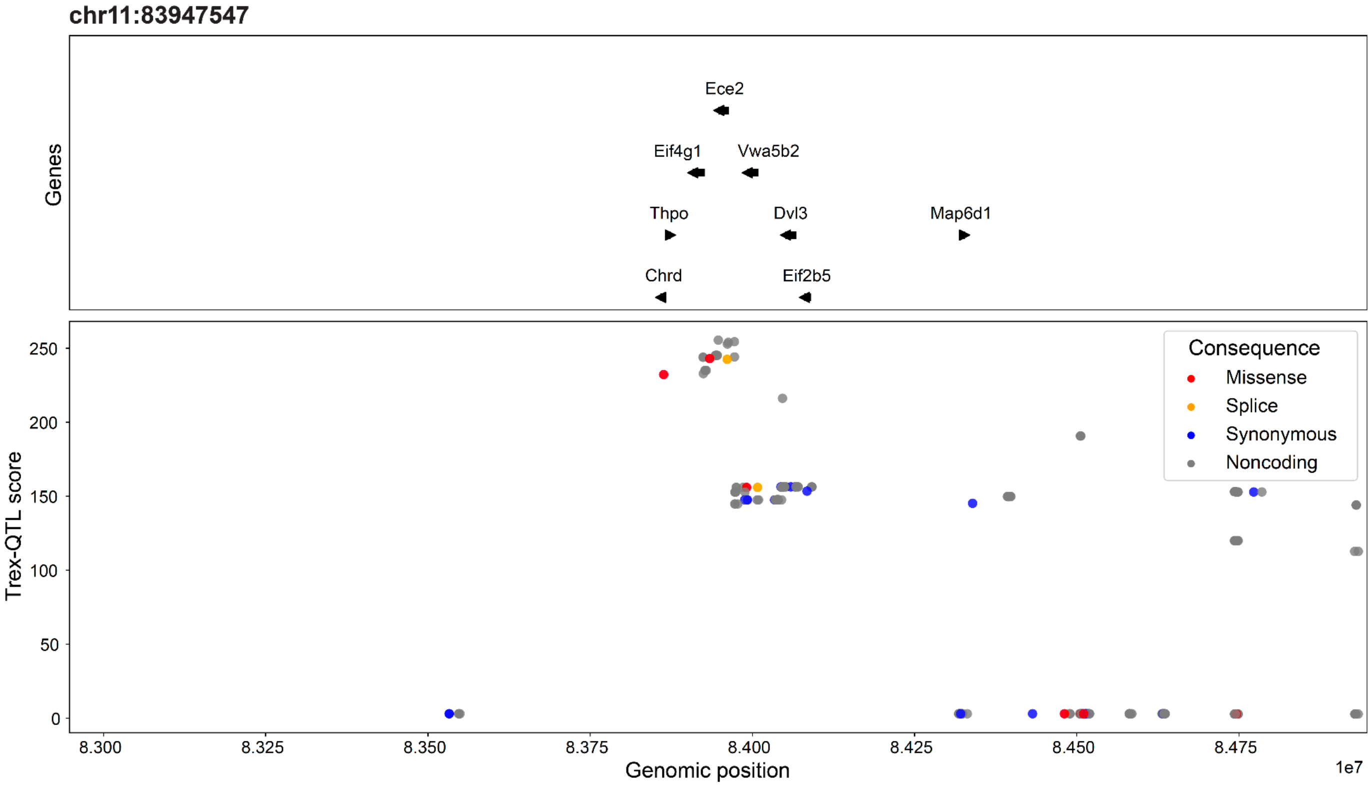


**Regional Trex-QTL association plots for top Trex-QTL loci.** Regional association plots showing *S* scores in the genomic region surrounding each lead trans-eQTL variant not shown in **Fig. 4a** and **Fig. 5a**. Variants are colored according to the most severe predicted functional consequence from the VEP: dark blue, synonymous; orange, splice variant; red, missense; and gray, noncoding. Only genes for which the lead *trans*-eQTL variant is also a significant *cis*-eQTL are shown. For the **chr9:111109123** locus, no genes are displayed because the lead variant was not a significant *cis*-eQTL for any nearby gene.

### Supplementary Fig. 8

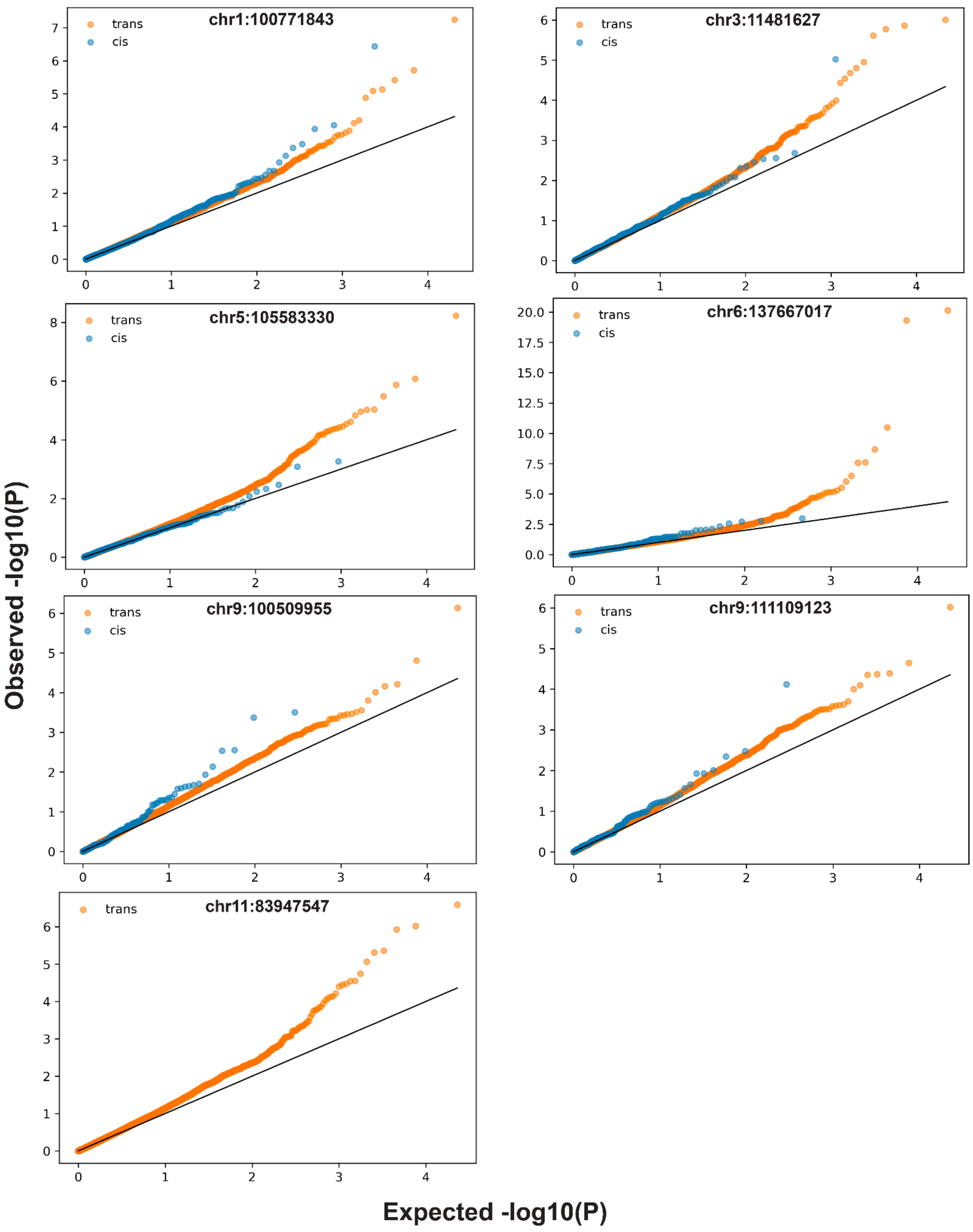

***Cis-trans* QQ plots for the 7 top *trans*-eQTL loci identified in the HS rat dataset.** For each candidate *trans*-eQTL, observed versus expected −log10(*P*) values from Matrix-eQTL association testing are shown separately for *cis* (blue) and *trans* genes (orange). *Cis* genes were defined as genes located on the same chromosome as the variant, whereas *trans* genes were located on all other chromosomes. Deviations above the diagonal indicate enrichment of association signals relative to the null expectation. Several loci exhibit widespread enrichment of *trans* associations across many genes, consistent with broad *trans*-regulatory effects, whereas others are characterized by a small number of stronger *trans* associations. As part of the Trex-QTL framework, the nearest *n* genes to each variant (*n* = 303, corresponding to the smallest number of tested genes on a single chromosome, Chromosome 11) were excluded from analysis to reduce residual *cis* effects while maintaining a constant number of tested genes across variants. Consequently, no *cis* genes are shown for the Chromosome 11 *trans*-eQTL locus.

### Supplementary Fig. 9

Lead variant: 1:100771843

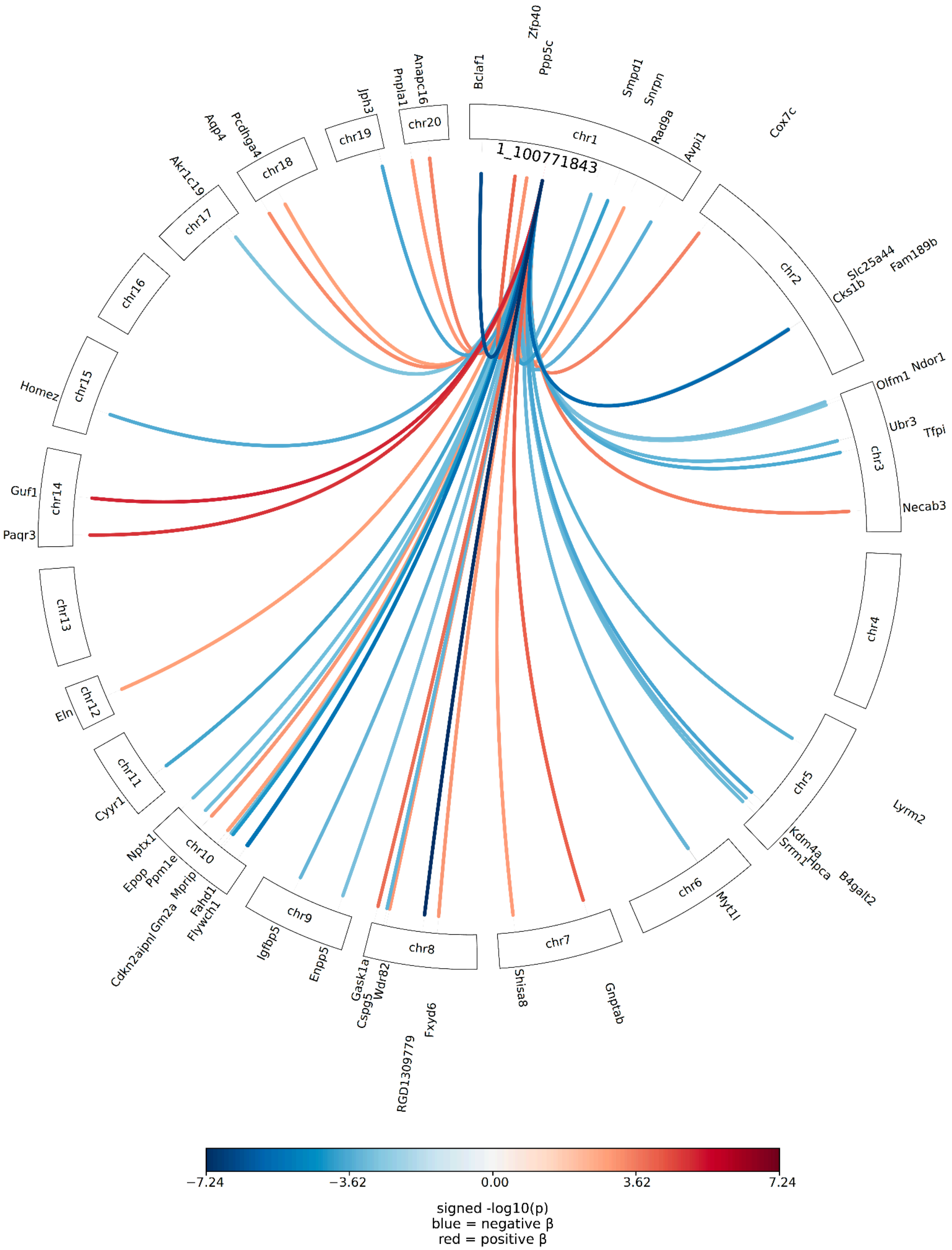

Lead variant: 3:11481627

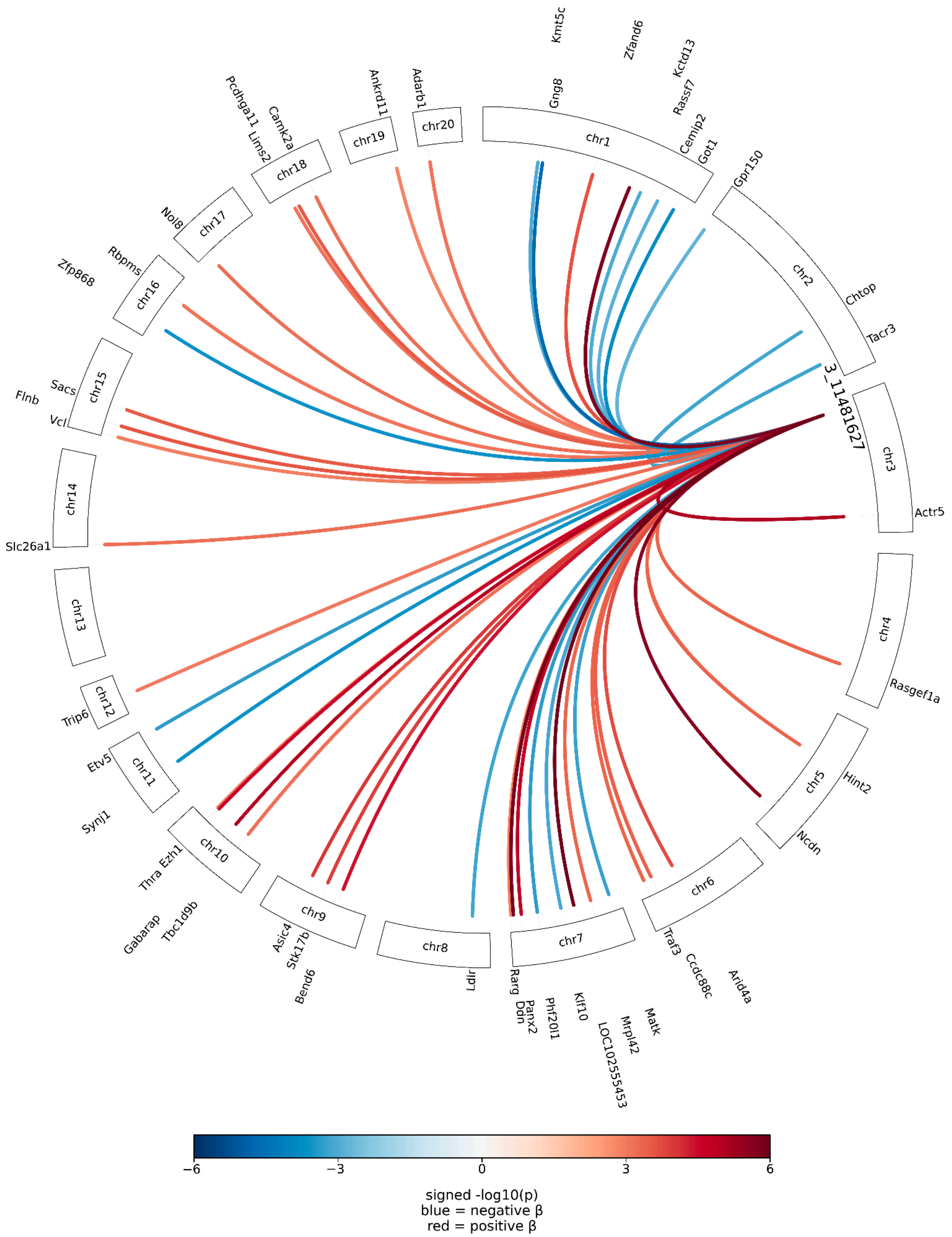

### Lead variant: 9:100509955

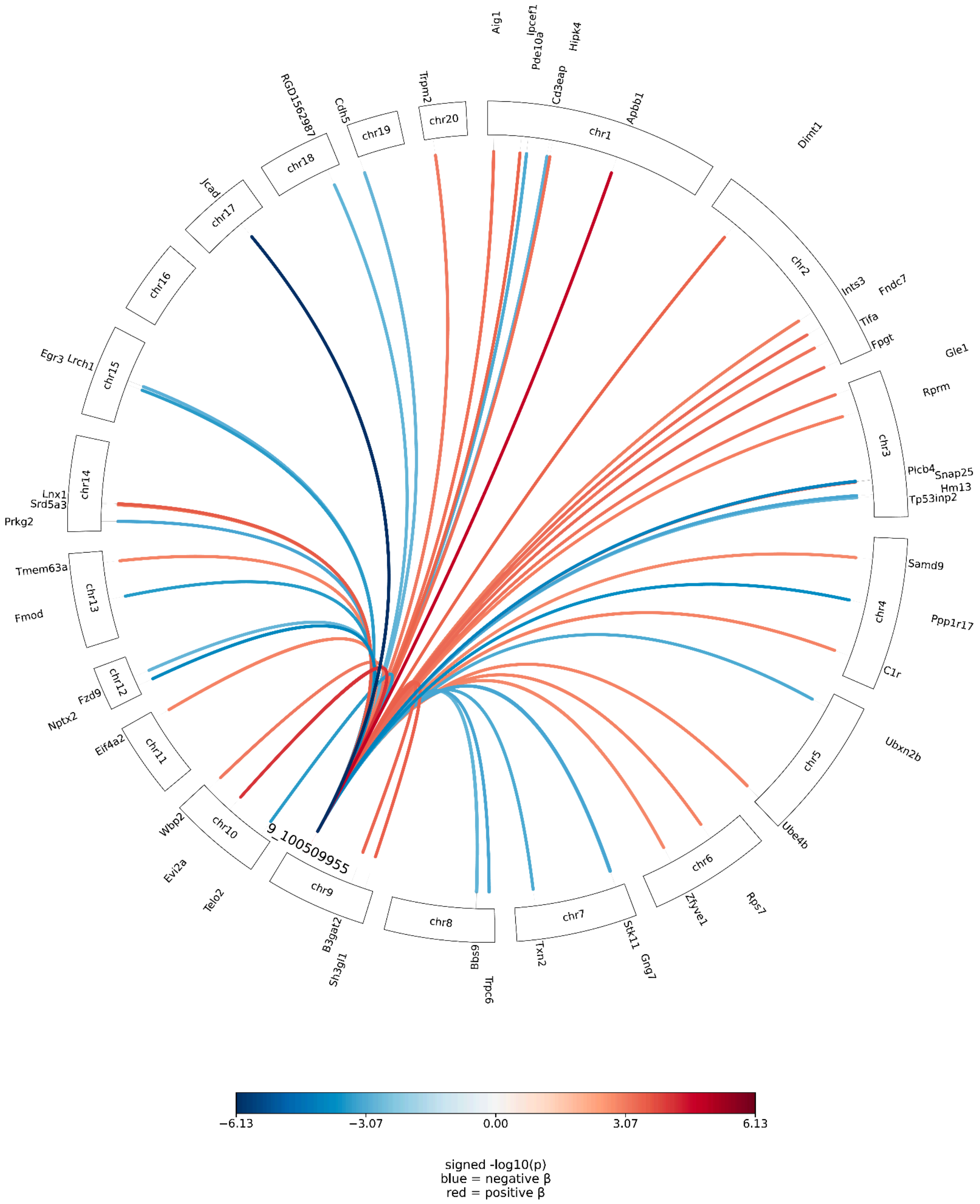

### Lead variant: 9:111109123

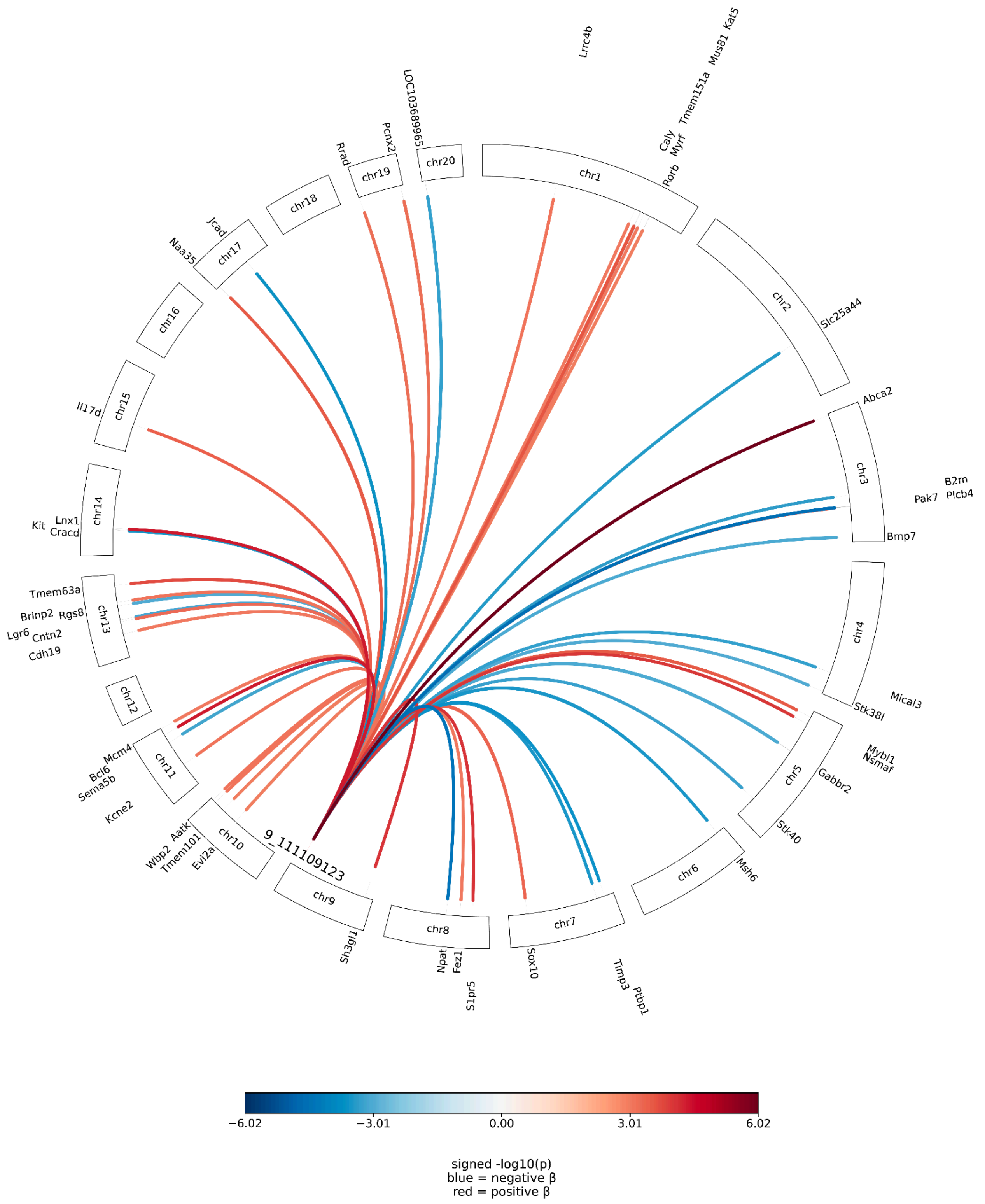

Lead variant: 11:83947547

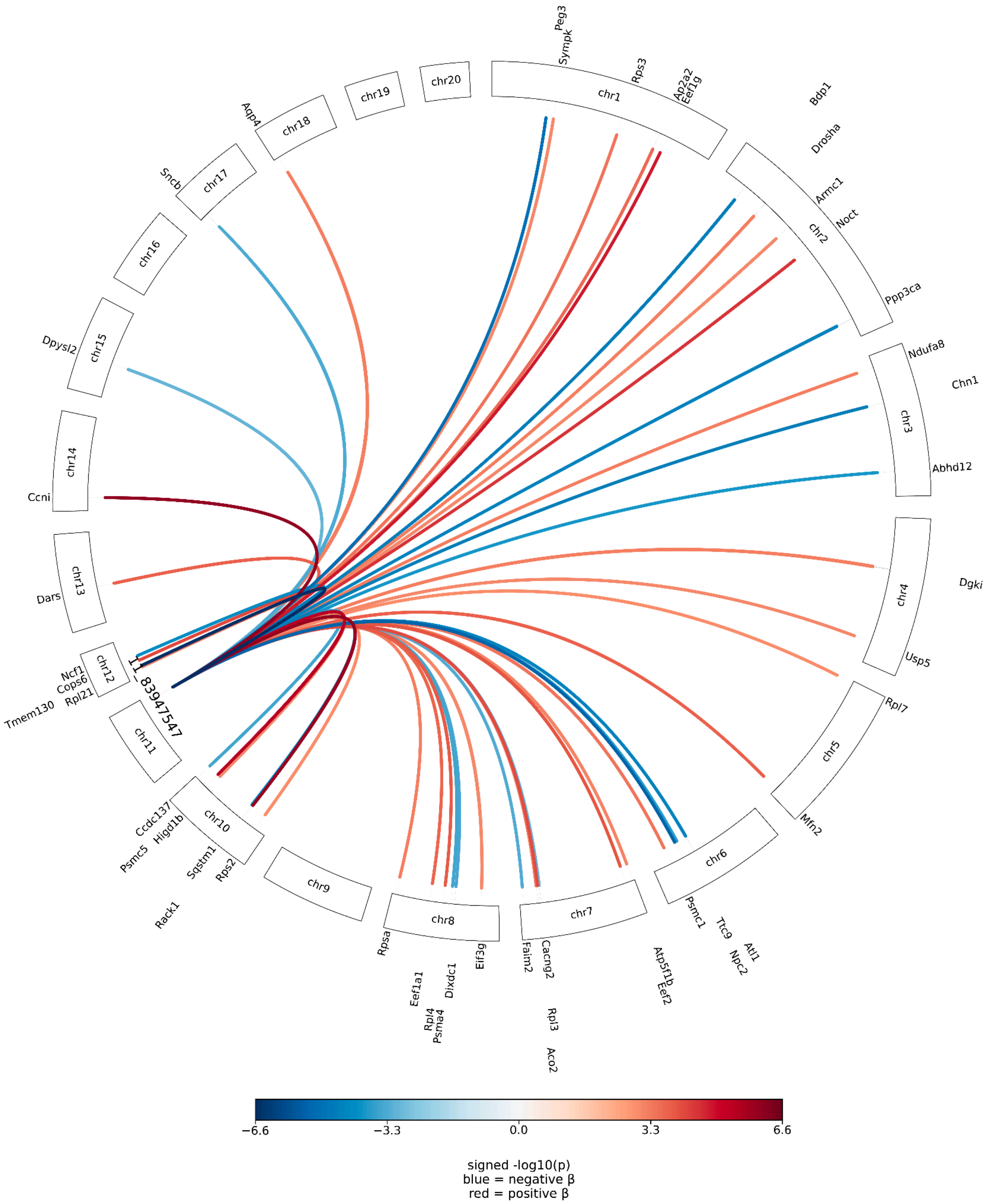


**Visualization of *trans* signals at significant Trex-QTL hotspots.** Genome-wide *trans*-eQTL networks for the significant Trex-QTL loci excluding the two loci shown in **Fig. 4f** and **Fig. 5d**. For each locus, the lead variant is connected to its target genes by links colored according to signed association strength (signed −log10(P)); blue indicates that the alternate allele is associated with decreased gene expression (negative $\beta$), whereas red indicates increased gene expression (positive $\beta$). Only the top 50 target genes for each locus are shown for clarity.

#### Supplementary Table and Dataset Legends

**Supplementary Table 1:** Trex-QTL association statistics for the initial set of 11,000 LD-pruned variants that passed preprocessing quality-control filters and were tested in the HS rat Trex-QTL analysis. Reported columns include the lead variant identifier, CPMA statistic and P-value, Trex-QTL likelihood ratio statistic (S), predicted proportion of target genes t (predicted_t), predicted mixture model parameter λ (predicted_L), Trex-QTL *P*-value, and Trex-QTL empirical *P*-value.

**Supplementary Table 2:** PLINK clumping results for significant Trex-QTL loci. For each lead Trex-QTL variant, the table lists all variants assigned to the corresponding LD clump, including both the lead variant and correlated variants. Clumping was performed after reintroducing pruned variants from candidate Trex-QTL regions.

**Supplementary Table 3:** Candidate Trex-QTL loci removed during quality-control filtering and the corresponding filtering criteria. Reported columns include the chromosome, genomic position, Trex-QTL likelihood ratio statistic (S), predicted proportion of target genes t, predicted mixture model parameter λ, and the reason each locus was removed.

**Supplementary Table 4:** Regional annotation of variants surrounding the Chromosome 6 Trex-QTL locus (chr6:137,667,017). Reported columns include chromosome, genomic position, Trex-QTL score, VEP consequence annotation, simplified functional consequence (Missense, Splice, Synonymous, or Noncoding), and the overlapping gene (if applicable).

**Supplementary Table 5:** Regional annotation of variants surrounding the chromosome 5 Trex-QTL locus (chr5:105,583,330). Reported columns include chromosome, genomic position, Trex-QTL score, VEP consequence annotation, simplified functional consequence (Missense, Splice, Synonymous, or Noncoding), and the overlapping gene (if applicable).

**Supplementary Dataset 1:** Full summary statistics for DGN analysis. Reported columns include the lead variant identifier, CPMA statistic and P-value, Trex-QTL likelihood ratio statistic (S), predicted proportion of target genes t (predicted_t), predicted mixture model parameter λ (predicted_L), Trex-QTL *P*-value, Trex-QTL empirical *P*-value, adjusted P-value, and whether the variant passes FDR 25%.
